# Role of Gasdermins in the Biogenesis of Apoptotic Cell–Derived Exosomes

**DOI:** 10.1101/2021.04.27.441709

**Authors:** Jaehark Hur, Yeon Ji Kim, Da Ae Choi, Dae Wook Kang, Jaeyoung Kim, Hyo Soon Yoo, Sk Abrar Shahriyar, Tamanna Mustajab, Dong Young Kim, Yong-Joon Chwae

**Affiliations:** Department of Microbiology, Ajou University School of Medicine, Suwon, Gyeonggi-do 16499, South Korea; Department of Biomedical Science, Graduate School of Ajou University, Suwon, Gyeonggi-do 16499, South Korea; Department of Medicine, Graduate School of Ajou University, Suwon, Gyeonggi-do 16499, South Korea; CK-Exogene Inc., Seoul 54853, South Korea; Department of Otolaryngology, Ajou University School of Medicine, Suwon, Gyeonggi-do 16499, South Korea

**Author notes:** These authors contributed equally to this work (Jaehark Hur, Yeon Ji Kim, Da Ae Choi, Dae Wook Kang, Jaeyoung Kim). **Address correspondence to:** Yong-Joon Chwae, Department of Microbiology, Ajou University School of Medicine, 164 World Cup Road, Yeongtong-gu, Suwon, Gyeonggi-do 443-380, South Korea.

## Abstract

The gasdermins, GSDMA, GSDMB, GSDMC, GSDMD, DFNA5, and DFNB59, are a family of pore-forming proteins that has recently been suggested to play a central role in the pyroptosis and the release of inflammatory cytokines. Here, we describe the novel roles of gasdermins in the biogenesis of apoptotic cell–derived exosomes. In apoptotic cells, GADMA, GSDMC, GSDMD, and DFNA5 increased the release of ApoExos, and both their full-length and cleaved forms were localized in the exosomal membrane. GSDMB and DFNB59, on the other hand, negatively affected the release of ApoExos. The caspase-mediated cleavage of gasdermins, especially DFNA5, is suggested to enable cytosolic Ca^2+^ to flow through endosomal pores and thus increase the biogenesis of ApoExos. In addition, the DFNA5-meidiated biogenesis of ApoExos depended on the ESCRT-III complex and endosomal recruitment of Ca^2+^-dependent proteins: annexins A2 and A7, the PEF domain family proteins sorcin and grancalcin, and the Bro1 domain protein HD-PTP. Therefore, we propose that the biogenesis of ApoExos begins when gasdermin-mediated endosomal pores increase cytosolic Ca^2+^, continues through the recruitment of annexin-sorcin/grancalcin-HD-PTP, and is completed when the ESCRT-III complex synthesizes intraluminal vesicles in the multivesicular bodies of dying cells. Finally, we found that Dfna5-bearing tumors released ApoExos to induce inflammatory responses in the *in vivo* 4T1 orthotropic model of breast cancer. The data presented in this study indicate that the switch from apoptosis to pyroptosis could drive the transfer of mass signals to nearby or distant living cells and tissues by way of extracellular vesicles, and that gasdermins play critical roles in that process.

## Introduction

Gasdermins, a family of pore-forming proteins, comprise six genes in humans: *GSDMA, GSDMB, GSDMC, GSDMD, DFNA5* (also known as *GSDME*), and *DFNB59* (also known as *PJVK*). All of them except DFNB59 consist of an N-terminal domain, a C-terminal domain, and a linker domain. Recent studies have demonstrated that the pore-forming activity of the N-terminal domains is prohibited by a structural auto-inhibitory function conferred by the C-terminal domains. The N-terminal domains can be activated by caspase-mediated cleavages in the linker domain, which results in the formation of membrane pores that evoke pyroptotosis and inflammation. GSDMD can be cleaved by proinflammatory caspases [caspases 1, 11 (in mice), or 1, 4, and 5 (in humans)], which allows it to act as the main executor of the inflammasome pathway; in that role, it facilitates the release of proinflammatory cytokines such as IL1β and IL18 and induces pyroptosis, thereby contributing to the innate immune response [1, 2, 3]. It has been recently noted that GSDMD can be activated by caspase 8 [4, 5] and the neutrophil elastase [6, 7, 8] suggesting that it could be implicated in other processes, such as the transition from apoptosis to pyroptosis and NETosis. Reportedly, DFNA5 can be cleaved by caspase 3 to carry out the pyroptotic transition in apoptotic cells [9, 10].

Although EVs have mostly been inspected in live cells during normal or stressed conditions, recent advances in EV research have highlighted EVs from dying cells [11]. Among them, apoptotic cell–derived exosomes (apoptotic exosomes: ApoExos), which originate from endosomes, are proposed to be secreted in a caspase 3- and 9-dependent manner. ApoExos have unique markers, including a sphingosine-1-phosphate receptor 1/3 (S1PR1/3), in addition to the known, exosome-specific tetraspanin CD63, which is involved in S1P/S1PR signaling and the subsequent endocytosis of the plasma membrane that occurs during ApoExo biogenesis [12, 13]. ApoExos have recently been shown to carry out a variety of functions, such as the induction of inflammation or an autoimmune response, the proliferation of tumors, and increasing cellular survival [11, 12].

Even though the synthesis of ApoExos is apoptotic caspase–dependent, no substrate has been found for the effector caspases that contribute to the biogenesis of ApoExos. In this research, we found that gasdermins play a key role in the maturation of MVBs during the biogenesis of ApoExos. Furthermore, we investigated the molecular mechanisms of the gasdermin-mediated maturation of MVBs and the biological significance of the gasdermins, focusing on DFNA5.

## Results

### The release of ApoExos can be regulated by gasdermin family genes

Cell lines showing higher expression of DFNA5 and GSDMD released more ApoExos although not in MCF7 cell line (Fig. S1). Moreover, overexpression of DFNA5 substantially increased the release of ApoExos, but DFNA5 knockout nearly completely blocked the release of ApoExos (Supplementary Fig. 2a and c). In addition, DFNA5 overexpression promoted pyroptotic cell death at an early phase of cell death; in contrast, DFNA5 knockout postponed cell death (Fig. S2b). Transiently expressed DFNA5 also increased the release of ApoExos in a dose-dependent manner (Fig. S3), and GSDMD knockout along with DFNA5 knockout reduced both the release of exosomes and cell death (Fig. S4).

Therefore, to further investigate the roles of gasdermins in the biogenesis of ApoExos, we asked whether the overexpression of each gasdermin would affect exosome release. To answer that question, we used treatment with staurosporine or TNFα and cycloheximide to induce apoptotic cell death in HeLa cells overexpressing gasdermins and then prepared ApoExos. We found that the overexpression of GSDMA, GSDMC, GSDMD, or DFNA5 considerably increased the release of ApoExos, as confirmed by western blotting and nanoparticle tracking analyses (NTAs). In contrast, the overexpression of GSDMB or DFNB59 did not increase and indeed significantly decreased the release of exosomes (Fig. 1a, b, d, and e, and S5). The overexpression of GSDMC, GSDMD, or DFNA5 augmented cell death, whereas overexpression of DFNB59 reduced cell death but only in staurosporine-treated cells (Fig. 1c and f). On the other hand, the overexpression of gasdermins in caspase 3 knockout cells eliminated the effects of GSDMA, GSDMC, and GSDMD on both the release of ApoExos and cell death, though DFNA5 still promoted the release of ApoExos and cell death. GSDMB and DFNB59 inhibited the release of ApoExos, consistent with the results from wild-type cells, and DFNB59 also prohibited pyroptotic cell death (Fig. 1g–1i). In 293T cells, DFNA5 increased the release of exosomes, but GSDMB and DFNB59 slightly suppressed the release of exosomes, analogous to the results in HeLa cells. GSDMA, GSDMC, and GSDMD showed no effects on the release of exosomes, which is inconsistent with the results in HeLa cells (Supplementary Fig. 6a and b). Meanwhile, GSDMA, GSDMC, GSDMD, and DFNA5 enhanced pyroptotic cell death, but DFNB59 partially blocked cell death (Fig. S6c).

**Fig. 1.**
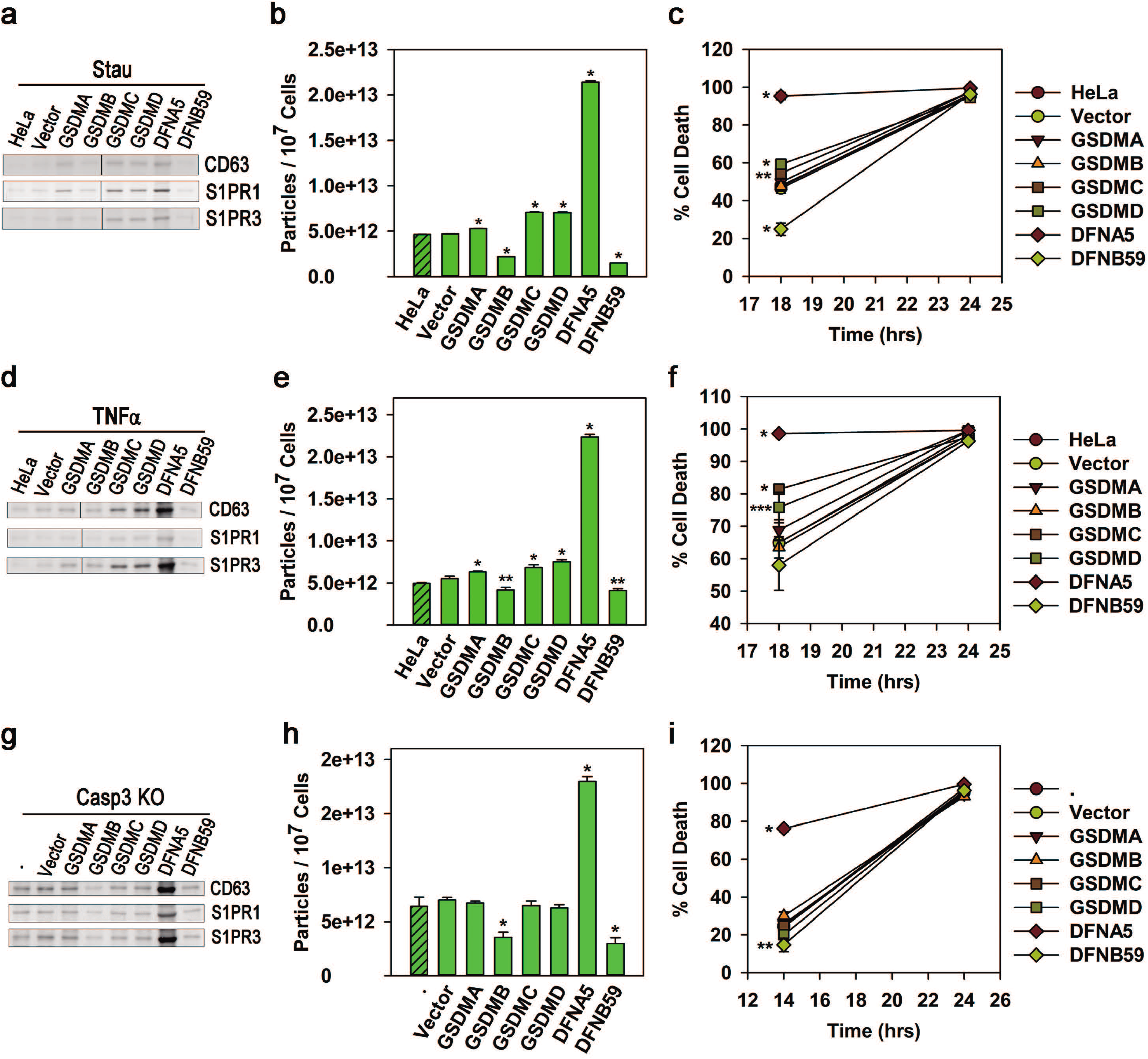
Gasdermins modulate the release of ApoExos. HeLa cells overexpressing a gasdermin protein and control cells were treated with staurosporine (1 μM) or TNFα (50 ng/ml) plus cycloheximide (25 μg/ml). HeLa cells depleted of caspase 3 and overexpressing a gasdermin were treated with staurosporine for 24 hr. **a, d,** and **g** ApoExos were prepared and western blotting was performed with equal volumes of exosomes for markers of ApoExos (CD63, S1PR1, or S1PR3). **b, e,** and **h** Equal volumes of the exosomal fractions were measured in a nanoparticle-tracking analysis (NTA). **c, f,** and **i** % cell death was calculated from the cells at the indicated times. * P < 0.001 and ** P < 0.01.

Collectively, our data demonstrate that DFNA5 promoted the release of ApoExos and enhanced pyroptotic cell death in all situations of apoptosis. On the other hand, GSDMB and DFNB59 played negative roles in the release of exosomes, and DFNB59 partially inhibited pyroptotic cell death. GSDMA-, GSDMC-, or GSDMD-mediated changes in exosome release and pyroptotic cell death depended on caspase 3 in a cell type–specific manner.

### Gasdermins are localized in the ApoExos in their full-length or cleaved form

Next, we examined where the gasdermins were localized on the membranes in which they subsequently made pores. Except for GSDMD, DFNA5, and DFNB59, the gasdermins were barely detectable in parental HeLa cells but observed by western blotting only in overexpressed cells (Fig. 2a). In overexpressed cells, GSDMA, GSDMC, GSDMD, and DFNA5 were detected in their full lengths and several cleaved forms in ApoExos. In particular, the full-length and cleaved forms of DFNA5 were detected in exosomes prepared from caspase 3 knockout cells. The full-length DFNB59 was also observed in the ApoExos. However, GSDMB could not be identified in the exosomes in any conditions (Fig. 2a). Thus, the gasdermins, including DFNA5, shown to promote the release of exosomes after apoptotic stimuli, need to be localized at the exosome in both their full-length and cleaved forms, and the negatively acting DFNB59 can be localized in the exosomes in its full-length form. To further confirm that point, we prepared exosomal membrane fractions and examined the localization of the gasdermins. We detected either the full-length or cleaved forms of GSDMA, GSDMC, GSDMD, DFNA5, and DFNB59 proteins in the exosomal membrane (Fig. S7). DFNA5 expression was also detected in our flow-cytometric analysis and in transmission EM images of exosomes (Fig. 2b and c).

**Fig. 2.**
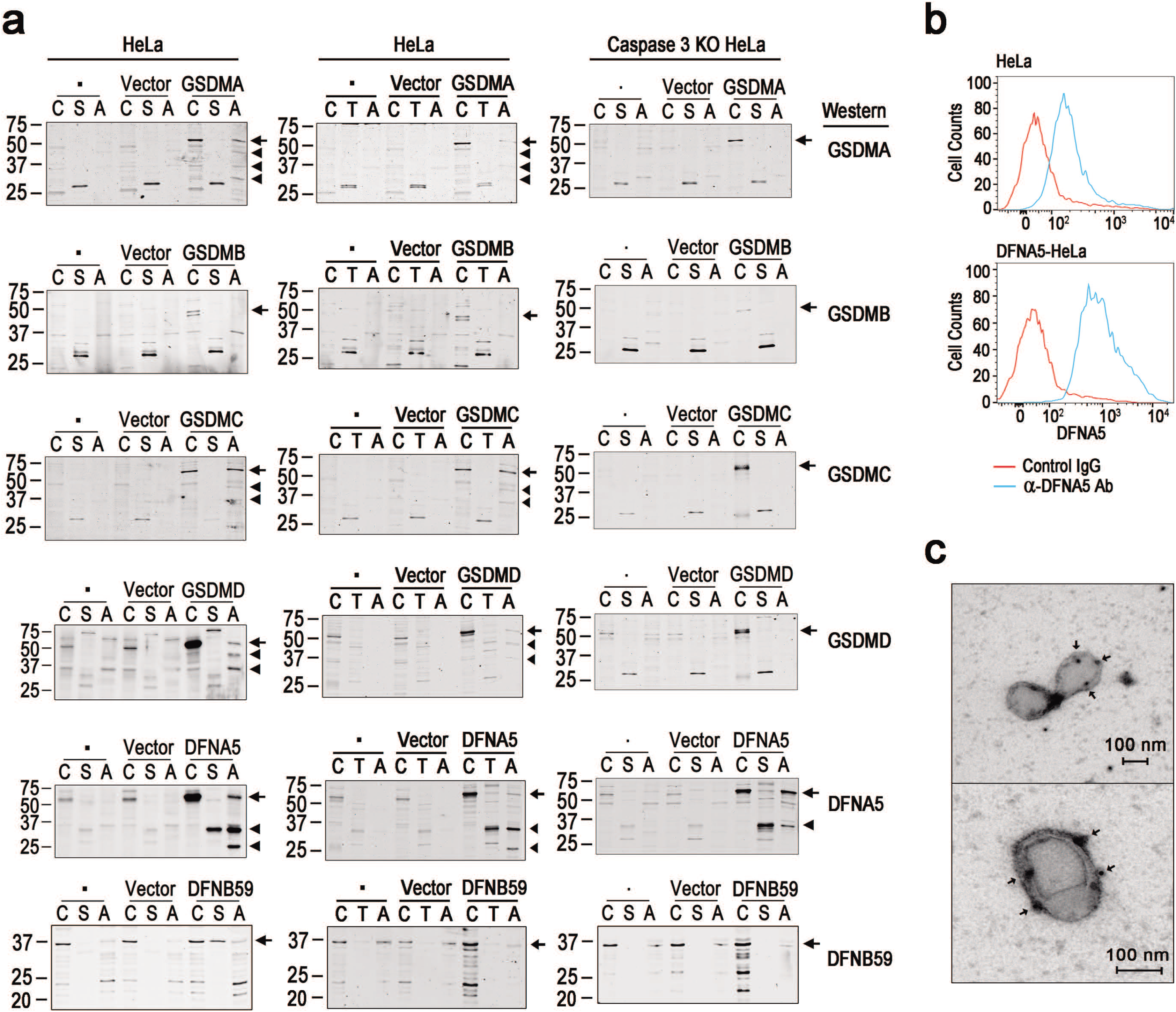
Gasdermins promoting the release of ApoExos are localized to those ApoExos in both their full-length and cleaved forms. **a** HeLa cells or caspase 3 knockout HeLa cells were made to overexpress a gasdermin (*GSDMA, GSDMB, GSDMC, GSDMD, DFNA5*, or *DFNB59*) or a transfected control vector (*Vector*). Those cells and parental HeLa cells were treated with staurosporine (1 μM) or TNFα (50 ng/ml) and cycloheximide (25 μg/ml) for 24 hr. The cell lysates and ApoExos were prepared, and western blotting was performed with equal amounts of protein for each gasdermin (C: cell lysates from non-treated control cells; S: cell lysates from staurosporine-treated cells; T: cell lysates from TNFα and cycloheximide–treated cells; A: lysates from the ApoExos). Arrows denote the full-length form of each gasdermin, and triangles denote the cleaved form of each gasdermin. **b** ApoExos from HeLa and DFNA5-overexpressing HeLa cells were coated onto aldehyde/sulfate beads and stained with anti-DFNA5 Ab (blue) or negative control Ab (red) to confirm the surface expression of DFNA5. **c** TEM images of the ApoExos stained with anti-DFNA5 Ab and a secondary Ab tagged with gold particles. Arrows indicate DFNA5s stained with gold particles.

### Caspase-mediated cleavage of gasdermins is required for maximum biogenesis of ApoExos

Next, we proposed that caspase-mediated cleavage and subsequent pore-formation were required for the gasdermin-mediated promotion of ApoExo biogenesis. To test that proposition, we replaced the Asp at the caspase-cleavage sites in DFNA5 and GSDMD with Glu to produce caspase-resistant mutants [9, 10, 14]. The cells expressing the mutant DFNA5 showed decreased release of exosomes and reduced cell death, but very interestingly, still induced the release of more exosomes and more cell death than in the control cells (Fig. 3b, c, and d). Consistently, the full-length DFNA5 mutants were able to be localized into the exosomes (Fig. 3a). However, in the cells expressing the GSDMD mutants, the release of exosomes was decreased to the levels of the control cells, and cell death decreased below the level of the control (Fig. 3f, g, and h). Neither the full-length nor cleaved form of the GSDMD mutants was localized in the exosomal fractions (Fig. 3e), implying that DFNA5 and GSDMD work in caspase-dependent manners; however, intact DFNA5 also plays a role in the biogenesis of ApoExos and pyroptosis. To further search for the gasdermin domains, we expressed non-pore-forming constructs of DFNA5 [9] (Fig. 4 a and b). As shown in figure 4c and d, the non-pore-forming fragments of DFNA5 were unable to increase the release of exosomes. In addition, the partial fragments interfered slightly with cell death (Fig. 4e). Next, we introduced N-terminal constructs, including the pore-forming segment, into the cells as a cumate-inducible system (Fig. 4f). The cumate-mediated induction of the pore-forming N-terminal fragment showed a similar or even modestly decreased release of ApoExos (Fig. 4g), and the early cell death patterns differed insignificantly from those of the non-treated controls (Fig. 4h). In contrast, the induction of full-length DFNA5 increased both ApoExo release and early pyroptotic cell death (Fig. S8).

**Fig. 3.**
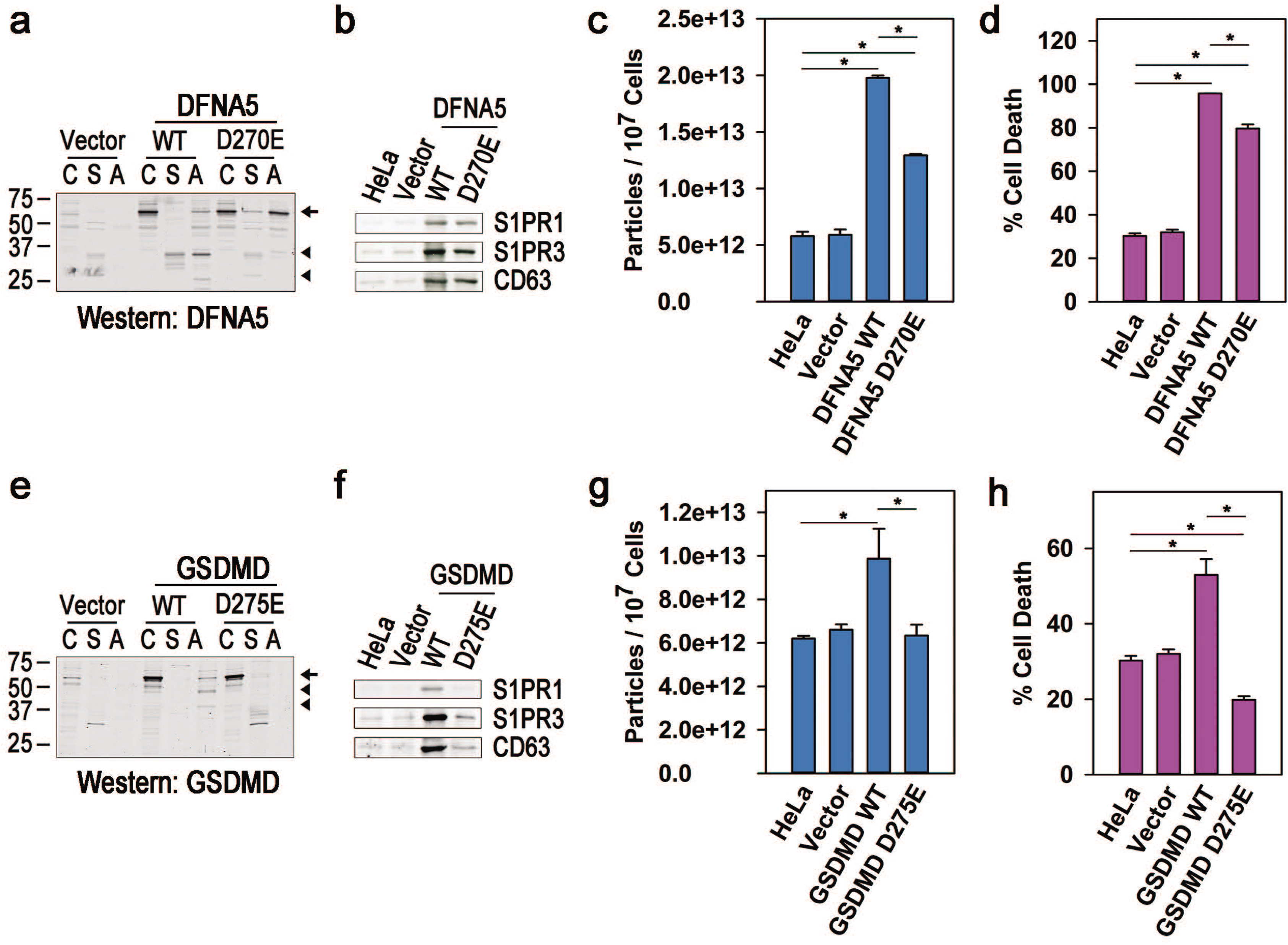
Mutation of caspase-cleavage sites in gasdermins partially blocked gasdermin-induced enhancement of apoptotic exosome release. **a** and **e** Control cells (*HeLa* or *Vector*) and cells overexpressing wild-type (*WT*) or mutant of either DFNA5 or GSDMD (*D270E or D275E*), were treated with staurosporine (1 μM) for 24 hr. the exosomal fractions were purified. Equal amounts of non-treated control cell lysates (*C*), staurosporine-treated cell lysates (*S*), and ApoExos (*A*) were analyzed by western blotting for DFNA5 or GSDMD. Arrows and triangles denote full-length and cleaved forms of gasdermin, respectively. **b, c, f,** and **g** Equal volumes of the exosomal fraction were western-blotted for markers of ApoExos (S1PR1, S1PR3, or CD63) and examined by NTA. **d** and **h** 12 hr after treatment, pyroptotic cell death was measured by the LDH assay. * P < 0.001 and ** P < 0.01.

**Fig. 4.**
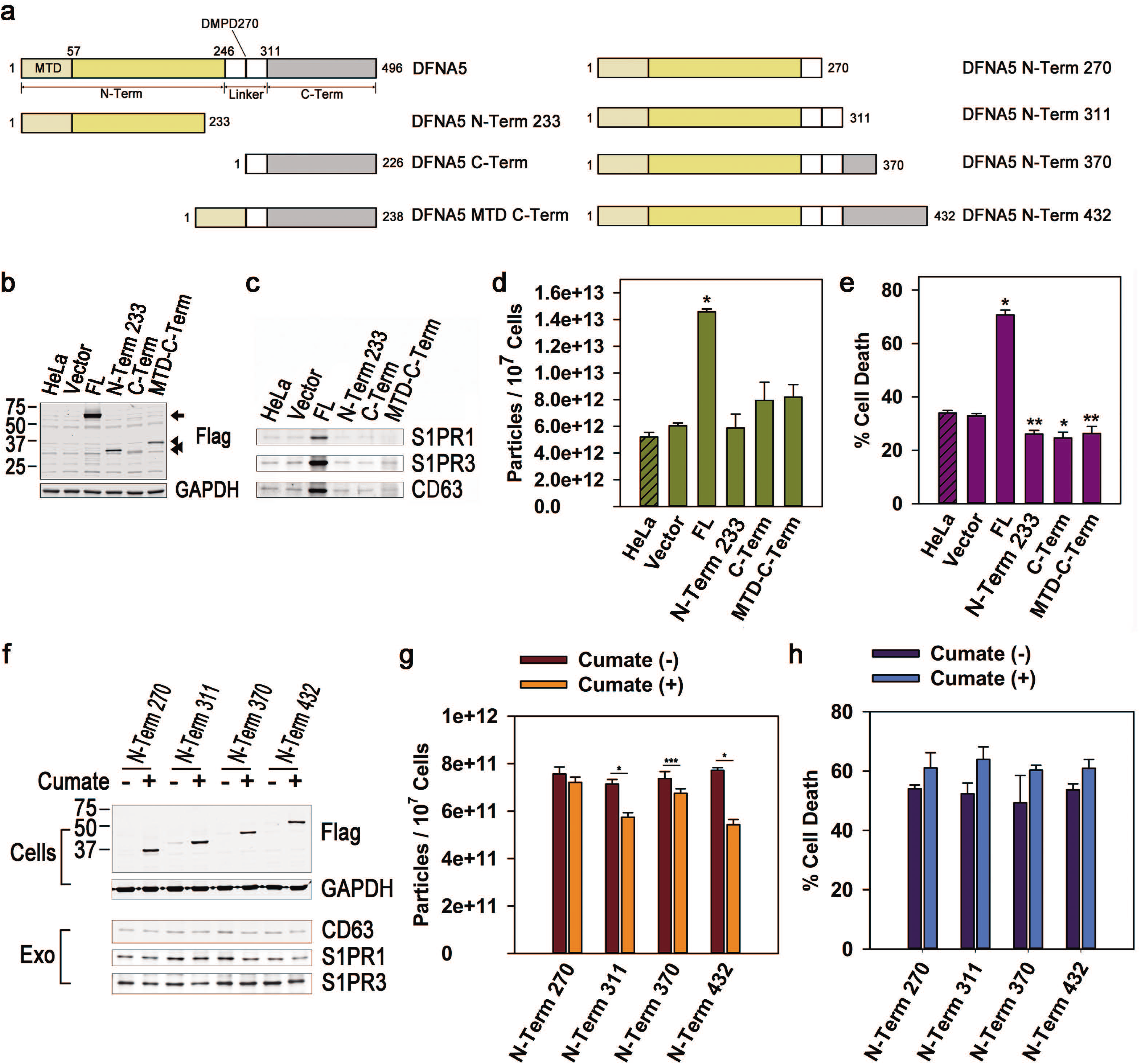
Full-length forms of DFNA5 protein are required for biogenesis of ApoExos. **a, left panel** Various constructs containing complete or partial cDNA sequences of DFNA5, as shown in the schematic diagram were infected into HeLa cells. **b** The cells were confirmed to express complete or partial cDNA of DFNA5 by western blotting with anti-Flag Ab. Arrows denote the full-length form, and triangles indicate fragmented forms of DFNA5. **c** and **d** Equal volumes of the ApoExos were analyzed by western blotting to examine the markers for ApoExos and by NTA to measure the concentration of exosomal particles. **e** pyroptotic cell death was detected by the LDH assay 12 hr after the staurosporine treatment,. **a, right panel** Partial or complete sequences of DFNA5 containing a pore-forming N-terminal segment were cloned into a cumate-inducible vector and stably expressed in HeLa cells. **f, upper panel** The cells were incubated in 20 μg/ml of cumate for 48 hr to induce the expression of the DFNA5 constructs, which was confirmed by western blotting. **f, lower panel** and **g** Then, the ApoExos were examined by western blotting and NTA. **h** Cell death was measured by the LDH assay around 12 hr after the treatment. * P < 0.001, ** P < 0.01, and ***P<0.5.

Taken together, these data suggest that caspase-mediated cleavage of gasdermins is important for the release of exosomes. Non-cleaved DFNA5 can increase the release of exosomes but not as efficiently as the cleaved form. This point is in accordance with a study reporting that gasdermins can be activated without caspase-mediated cleavage [15]. Intriguingly, the pore-forming N-terminal domain did not increase the release of the exosomes, indicating that the C-terminal auto-inhibitory domain is also essential for exosome biogenesis.

### DFNA5-mediated increase in the release of ApoExos is mediated by ESCRT complex activation

When we compared cells depleted of DFNA5 with parental cells in apoptotic conditions under a confocal microscope, the DFNA5 knockout cells showed marked reduction in the size of the intracellular vesicles co-localized with CD63, although early apoptotic changes, including the formation of plasma membrane spikes and the movement of CD63 toward them [13], differed little between the two groups (Fig. 5a and S9). Those differences could also be seen in transmission EM. Apoptotic cells were characterized by large intracellular vesicles, possibly MVBs, which were barely detectable in the DFNA5 knockout cells (Fig. 5b). Thus, the number of intracellular vesicles larger than 1 μm in diameter per cell was significantly reduced in the DFNA5 knockout cells (Fig. 5c). When cells expressing DFNA5 with an internal or C-terminal Flag tag (Fig. S10), were treated with staurosporine, the DFNA5 was cleaved into N-terminal (30 kDa) and C-terminal domains (20 kDa) in a time-dependent manner, and the C-terminal domain was rapidly degraded (Fig. 5d). In confocal studies, however, DFNA5 proteins internally fused with mCherry (similar to GSDMD internally fused with a fluorescence protein at the N-terminal region of the caspase-cleavage site in the linker domain [16] (Fig. S10)) were exactly co-localized with CD63 as aggregates of large intracellular vesicles (Fig 5e). Therefore, the full-length and cleaved forms of DFNA5 are localized to MVBs during apoptotic processes and appear to be implicated in their maturation.

**Fig. 5.**
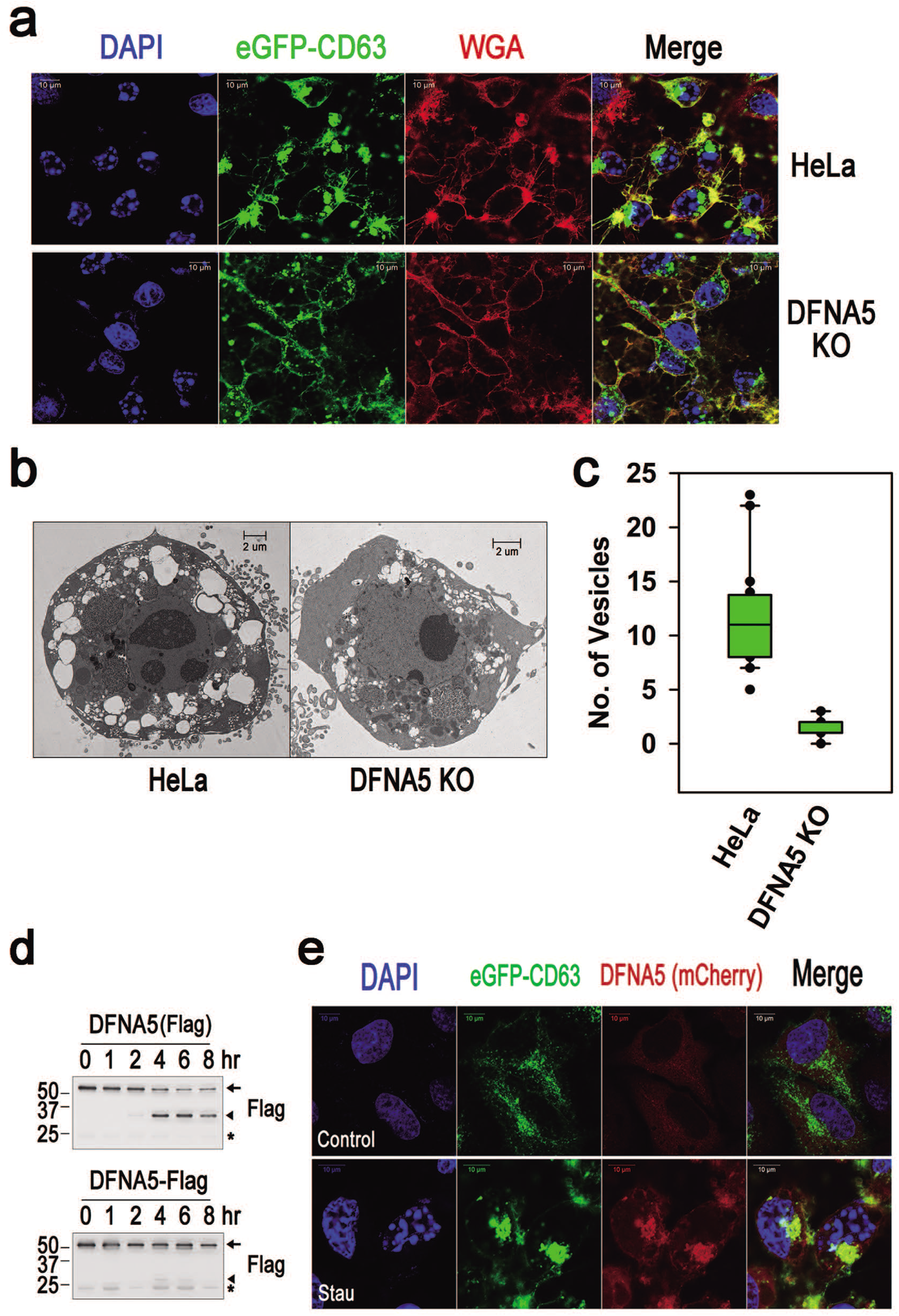
DFNA5 is involved in the maturation of multivesicular endosomes during apoptotic exosome biogenesis. **a** HeLa cells (*HeLa*) and HeLa cells depleted of DFNA5 (*DFNA5 KO*) expressing eGFP-CD63 were treated with staurosporine (1 μM) for 5 hr. The cells were then stained with wheat germ agglutinin conjugated with Alexa Fluor 594 (*WGA*) to visualize the plasma membrane. **b** and **c** HeLa cells and cells deprived of DFNA5 were treated with staurosporine (1 μM) for 7 hr, and then TEM images were made. One representative image from each group is shown. Intracellular vesicles more than 1 μm in diameter were counted in 20 individual cells from each group. **d** HeLa cells expressing DFNA5 tagged with internal or C-terminal Flag [*DFNA5(Flag*) or *DFNA5-Flag*] were treated with staurosporine. DFNA5 was immunoprecipitated and western blotted for Flag. Arrows and triangles denote the full-length and cleaved forms of DFNA5, respectively. Asterisks indicate immunoglobulin light chains. **e** HeLa cells expressing eGFP-CD63 and DFNA5 with internal mCherry [DFNA5(*mCherry*)] were treated with DMSO or staurosporine (1 μM) for 5 hr. Confocal images are shown.

Next, we investigated whether the DFNA5-mediated maturation of MVBs depended on ESCRT complexes. To do that, we examined how a dominant-negative mutant of VPS4B, VPS4B E235Q [17], and deleted mutants of CHMP2A and CHMP3 at the C-terminal region, which functioned as dominant-negative forms of ESCRT complexes [18], affected the DFNA5-mediated increase of ApoExos. The expression of the dominant-negative mutants of VPS4B, CHMP2A, or CHMP3 (Fig. 6a and c) diminished the release of ApoExos (Fig. 6a-d). Furthermore, although the overexpression of DFNA5 in cells expressing dominant-negative mutants of ESCRT complex members (Fig 6e) slightly increased the exosome release, it did not reach the levels of other cells overexpressing DFNA5 (Fig 6e and f). As shown by confocal microscopy, DFNA5 with internal eGFP was co-localized with members of the ESCRT-III complex, CHMP2A or CHMP4B in the intracellular MVBs (Fig. 6g and S11). In addition, immunoprecipitation experiments demonstrated that full-length and N-terminal fragments of DFNA5 were able to bind to CHMP4B in apoptotic cells (Fig. 6h). Interestingly, the interaction between DFNA5 and CHMP4B depended on the presence of Ca^2+^ (Fig. S12). Thus, DFNA5-mediated mechanisms are used in the steps of MVB maturation through Ca^2+^-dependent ESCRT activation.

**Fig. 6.**
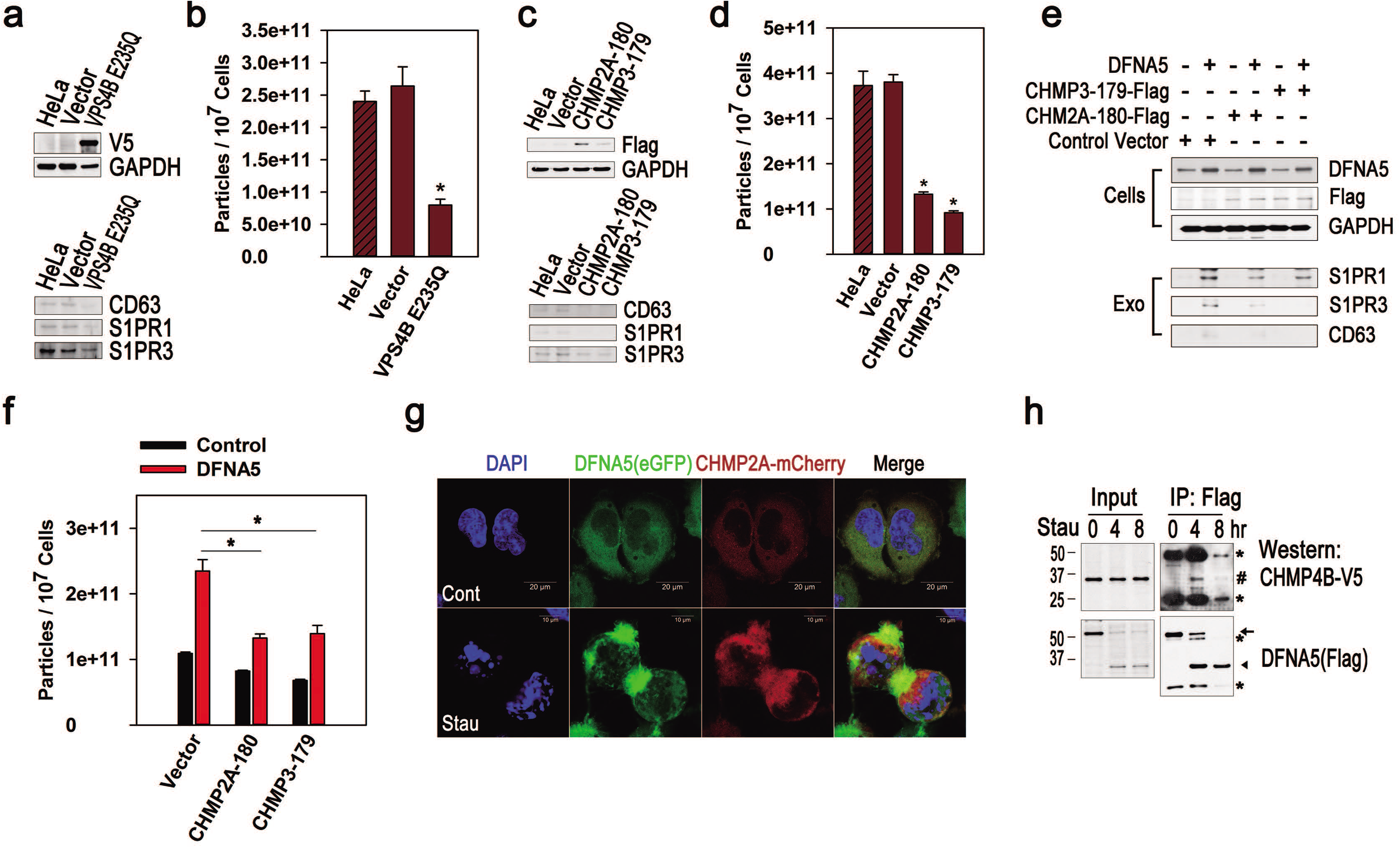
DFNA5-mediated promotion of apoptotic exosome biogenesis is executed through the activation of ESCRT complexes. **a** and **b** Control cells and cells overexpressing mutant VPS4B (*VPS4B E235Q*) (**a, upper panel**) were treated with staurosporine (1 μM) for 24 hr. ApoExos were analyzed by western blotting (**a, lower panel**) and NTA (**b**). **c** and **d** Cells overexpressing partial CHMP2A or CHMP3 (*CHMP2A-180* or *CHMP3-179*) (**c, upper panel**) were incubated in staurosporine for 24 hr, and the ApoExos were analyzed by western blotting (**c, lower panel**) and NTA (**d**). **e** and **f** Control cells and cells expressing mutants of CHMP2A or CHMP3 were infected with an empty vector or DFNA5 cDNA, and their expression of DFNA5 or the mutant CHMP2A or CHMP3 was confirmed by western blotting (**e, upper panel**). Exosomes from cells treated with staurosporine for 24 hr were examined by western blotting (**e, lower panel**) and NTA (**f**). **g** Cells overexpressing DFNA5 with internal eGFP and CHMP2A fused with mCherry were treated with staurosporine or solvent for 5 hrs. Confocal images are shown. **h** Cells were incubated with staurosporine for the indicated times. DFNA5 was immunoprecipitated with anti-Flag Ab, and western blotting was performed using anti-V5 or anti-Flag in the post-nuclear lysates containing Ca^2+^. Arrows and triangles denote the full-length and cleaved N-terminal fragments of DFNA5, respectively. # denotes CHMP4B tagged with V5, and asterisks denote heavy and light chains of Ig. *P < 0.001 (**b, d,** and **f**).

### Annexins are implicated in the DFNA5-mediated increase in ApoExo biogenesis

Given the Ca^2+^-dependent interaction between DFNA5 and the ESCRT-III complex, it is reasonable to think that DFNA5 forms pores in the endosome through which Ca^2+^ migrates to increase the regional cytosolic Ca^2+^ concentration, which in turn promotes the recruitment ESCRT-III [19]. To examine that notion, we tested the role of Ca^2+^ in DFNA5-mediated ApoExo production. Intracellular free Ca^2+^ was gradually decreased in both wild-type and DFNA5-depleted cells during early apoptotic cell death but transiently increased in cells overexpressing DFNA5 (Fig. S13). Co-treatment with an intracellular calcium chelator, BAPTA-AM, during staurosporine application blocked almost all of the ApoExo release in all the cells. However, co-treatment with a calcium ionophore, A23187, caused a considerable increase in ApoExo release in wild-type and DFNA5-depleted cells (Fig. 7a and b), however eliminated ApoExo release in DFNA5 overexpressing cells, indicating that Ca^2+^ is essential for ApoExo release and the fine control of regional Ca^2+^ mediated by the DFNA5 pores is required for the biogenesis of ApoExos.

**Fig. 7.**
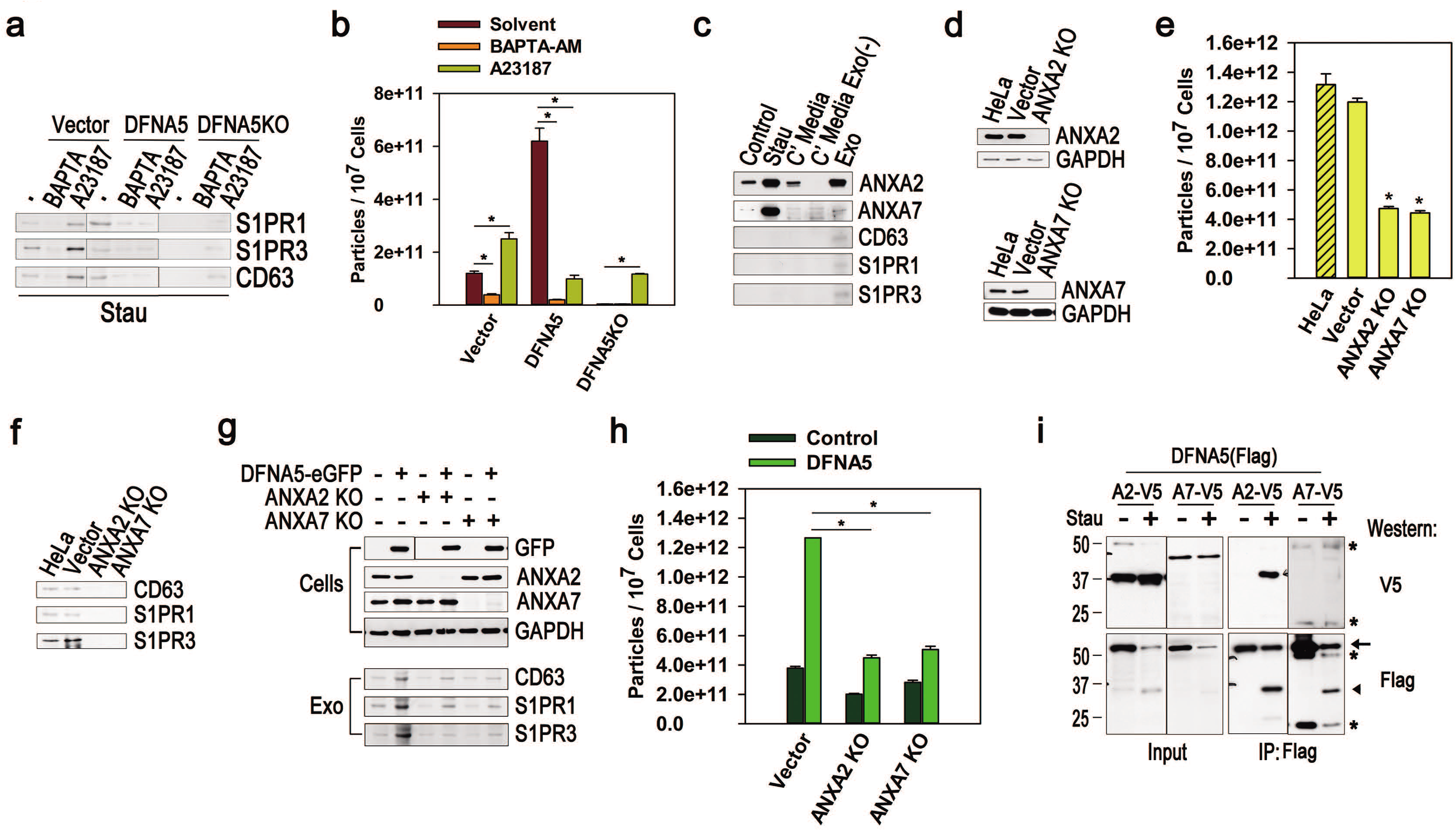
Ca^2+^/anionic phospholipid-binding proteins, annexins are implicated in DFNA5-mediated increase in apoptotic exosome biogenesis. **a** and **b** Vector-infected HeLa cells, cells overexpressing DFNA5, and DFNA5 KO cells were treated with staurosporine (1 μM) for 24 hr with solvent, BAPTA-AM (50 μM), or A23187 (1 μg/ml), and then ApoExos were prepared from the conditioned media. The exosomes were analyzed by western blotting (**a**) and NTAs (**b**). **c** The total cellular lysate (*Stau*), concentrated conditioned media (*C’ Media*), conditioned media depleted in the exosomal fractions [*C’ Media Exo(-)*], and ApoExos (*Exo*) were prepared from HeLa cells treated with staurosporine for 24 hr and compared with control cell lysate (*Control*) by western blotting with the indicated Abs. **d** Depletion of ANXA2 or ANXA7 was confirmed by western blotting in ANXA2 and ANXA7 knockout cells, which were treated with staurosporine for 24 hr. **e** and **f** The ApoExos were purified and analyzed by NTAs and western blotting. **g, upper panel** Wild-type cells, ANXA2-depeleted, and ANXA7-depleted cells were infected with DFNA5-eGFP cDNA or an empty vector, and their lysates were western-blotted with the indicated Abs. **g, lower panel** and **h** The ApoExos were western-blotted for CD63, S1PR1, or S1PR3, and measured by NTAs. **i** Cells expressing DFNA5(Flag) and ANXA2-V5 were incubated with a solvent control or staurosporine for 4 hr, and then their lysates were immunoprecipitated with anti-Flag Ab (*IP: Flag*) and western-blotted with anti-V5 Ab or anti-Flag Ab. One-hundredth volume of the lysates was western blotted with anti-V5 Ab or anti-Flag Ab (*Input*). Asterisks indicate Ig heavy and light chains. Arrow and triangle indicate full-length and the N-terminal domain of DFNA5, respectively. *P < 0.001 (**b, e,** and **h**).

Next, we looked for the link connecting DFNA5-mediated regional Ca^2+^ increase with recruitment of the ESCRT-III complex. To do that, we explored the effects of annexin A2 (ANXA2) and annexin A7 (ANXA7), which are Ca^2+^- and anionic phospholipid–binding proteins, on ApoExo release because ANXA2 and ANXA7 were previously shown to be associated with the recruitment of the ESCRT-III complex in plasma membrane repair [20, 21]. We prepared post-nuclear cell lysates, conditioned media, conditioned media depleted of their exosomal fractions, and ApoExos from apoptotic cells and then compared the expression of ANXA2 or ANXA7 among the groups. We found that both ANXA2 and ANXA7 were concentrated in the lysates of dead cells suggesting that they were located in the membrane fraction. In addition, ANXA2 was concentrated in the exosomal fractions, but ANXA7 was not (Fig. 7c). Depletion of either ANXA2 or ANXA7 (Fig. 7d) reduced the release of ApoExos (Fig. 7e and f). Furthermore, DFNA5 overexpression in ANXA2 or ANXA7 knockout cells partially restored ApoExo release but showed markedly reduced release compared with DFNA5 overexpression in wild-type cells (Fig. 7g and h). Immunoprecipitation of either ANXA2 or ANXA7 co-immunoprecipitated N-terminal DFNA5 domain in apoptotic cells (Fig. S14), and ANXA2 or ANXA7 was detected in the immunoprecipitates of DFNA5 (Fig. 7i). In the confocal images, ANXA2 or ANXA7 was co-localized with DFNA5 at the MVBs of apoptotic cells (Fig. S15). However, overexpression of ANXA2 or ANXA7 partially blocked ApoExo release (Fig. S16), indicating that DFNA5-induced endosomal pores invoke a cytosolic increase in Ca^2+^, resulting in the recruitment of annexins to the endosomal membrane, though the maintenance of an adequate amount of annexin is needed during the process.

### Sorcin/grancalcin-HD-PTP recruits ESCRT-III to the MVBs, resulting in the biogenesis of ApoExos

Based on a report demonstrating that Ca^2+^-activated ANXA7 forms a complex with apoptosis linked gene 2 (ALG2) and ALG2-interacting protein X (ALIX) to guide the ESCRT-III complex to the site of damage on a membrane [21], we wondered whether penta-EF hand (PEF) Ca^2+^-binding proteins such as ALG2, peflin, sorcin, and grancalcin [22] could manage the release of ApoExos. As shown in Figure 8, overexpression of sorcin or grancalcin increased the release of ApoExos; in contrast, the overexpression of ALG2 or peflin eliminated the release of ApoExos (Fig. 8a and b). In addition, knockout of sorcin or grancalcin blocked the release of ApoExos (Fig. S17). In agreement with those results, sorcin and grancalcin were co-localized with DFNA5 in the MVBs of apoptotic cells, but ALG2 and peflin were not (Fig. 8c). Sorcin and grancalcin also co-immunoprecipitated with ANXA2, ANXA7 and CHMP4B in apoptotic cells (Fig. 8d). Next, we investigated the effect of two Bro1 domain–containing proteins, ALIX and HD-PTP (His domain-containing protein tyrosine phosphatase), on the release of ApoExos. The Bro1 domain has been shown to bind to subunits of the ESCRT-III complex [23]. ALIX knockout did not decrease the release of ApoExos but instead increased it slightly (Fig. S18), consistent with a previous report [13]. The HD-PTP knockout cells reduced the release of ApoExos (Fig. 8e and f). HD-PTP in apoptotic cells was co-localized with DFNA5 in the cytosolic MVBs (Fig. 8g). Subcellular fractionation studies revealed that cleaved DFNA5 and the full-length form migrated into both the plasma membrane–containing heavy membrane fraction (HMF) and the MVB-containing light membrane fraction (LMF) in apoptotic cells, although it existed mainly in the cytosolic fraction in the resting cells. In the LMFs, the cleaved DFNA5 was co-localized with ANXA2, ANXA7, sorcin, grancalcin, HD-PTP, and CHMP4B (Fig. S19).

**Fig. 8.**
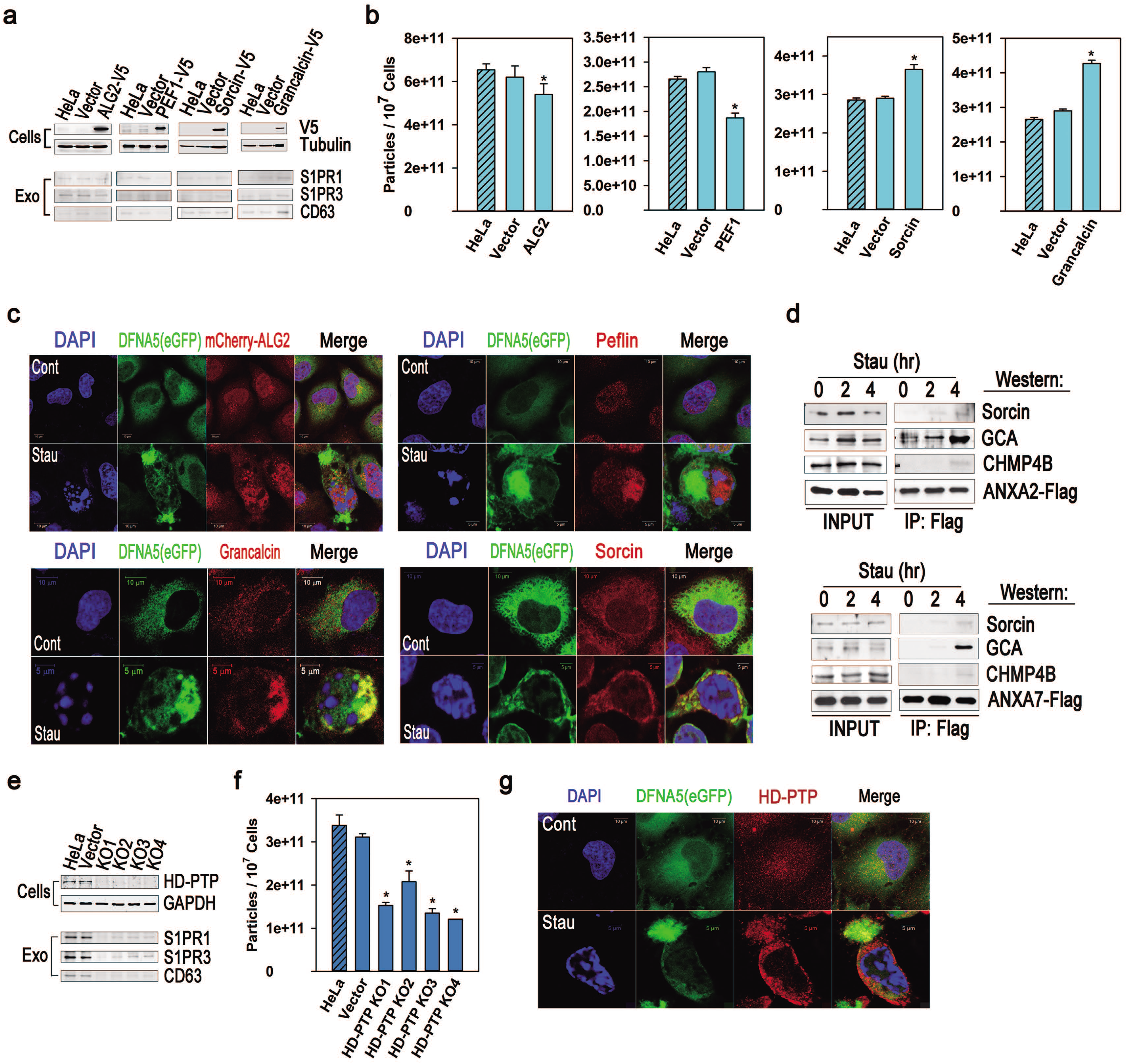
Sorcin/grancalcin-HD-PTP-ESCRT-III axis is associated with the maturation of MVBs in the biogenesis of ApoExos. **a, upper panel** Genes of the PEF family were overexpressed, and their expression was confirmed by western blotting. **a, lower panel** and **b** ApoExos were measured by western blotting and NTAs. **c** Cells were treated with solvent or staurosporine for 4 hr, after which confocal images were made. **d** Cells expressing ANXA2-Flag or ANXA7-Flag were lysed after staurosporine treatment (1 μM) for the indicated times and immunoprecipitated with anti-Flag Ab. The immunoprecipitates were western-blotted with the indicated Abs (*IP: Flag*). 1/100 volume of the lysate was western-blotted with the indicated Abs (*INPUT*). **e, upper panel** HeLa cells depleted of HD-PTP by the CRISPR/Cas9 system and control cells were western-blotted for HD-PTP. **e, lower panel** and **f** Then, ApoExos were analyzed by western blotting and by NTAs. **g** Cells expressing DFNA5(eGFP) were treated with staurosporine for 4 hr and stained for HD-PTP, and then confocal images were taken. *P < 0.001(**a** and **f**).

Taken together, our results suggest that sorcin and grancalcin, activated by an increase in cytosolic Ca^2+^ bind to annexins, and form a complex with HD-PTP, which enhances the assembly of the ESCRT-III complex through its Bro1 domain to contribute to the formation of ILVs. Thus, the biogenesis of ApoExos requires gasdermin-mediated calcium flow and the subsequent formation of an annexin-sorcin/grancalcin-HD-PTP-ESCRT-III axis.

### DFNA5 overexpression enhances levels of ApoExos and inflammation in the 4T1 orthotropic breast cancer model

To confirm the biological significance of gasdermin-mediated ApoExo release, we investigated the effect of Dfna5 overexpression in the *in vivo* 4T1 orthotropic breast cancer model [24]. 4T1 cells stably-expressing Dfna5 increased ApoExo release (Fig. S20a and b). The cells were injected into the mammary fat pads of BALB/c mice. When the tumors reached 1,000 mm^3^, the mice were injected with the anticancer drugs cyclophosphamide and doxorubicin or PBS, and then plasma exosomes and inflammatory mediators in the liver tissue were analyzed. As shown in Supplementary Figure 20c, the growing tumors showed no influence on the total amount of exosome compared with control mice without tumors, but anticancer drug treatments increased the absolute amount of plasma exosomes, even though the control tumors and Dfna5-overexpressing tumors did not differ. In contrast, ApoExos certainly increased in Dfna5-expressing tumors treated with anticancer drugs (Fig. S20d). Furthermore, the mice with Dfna5-expressing tumors induced more inflammatory mediators, including Il1b and Cox2, than found in the control groups (Fig. S20e and f). Thus, Dfna5 in mouse tumor tissues enhanced the release of ApoExos, which subsequently induced inflammatory responses in tumor-bearing mice, supporting previous findings on the inflammatory role of ApoExos [13, 25, 26].

## Discussion

The proposed mechanisms for the gasdermin-mediated biogenesis of ApoExos supported by the data presented here are summarized in a schematic illustration (Fig. S21).

GSDMB and DFNB59 blocked the release of ApoExos, and DFNB59 also prohibited early cell death in all conditions (Fig. 1). Thus, GSDMB and DFNB59 probably prevent pore formation in the membrane via other gasdermins, which was an unexpected result because the overexpression of the GSDMB N-terminal domain induced cell death, and DFNB59 has an N-terminal domain similar to that of DFNA5 [15, 27]. Possibly, a single nucleotide polymorphism found in GSDMB or a mutation of DFNB59 would alleviate the inhibitory effect of both proteins on membrane-pore formation, which would contribute to inflammation and immune responses to autoantigens in association with asthma and autoimmune diseases or neurosensory hearing loss, respectively [28].

So far, it remains unclear how GSDMA and GSDMC could be processed to generate N-terminal domains [29]. From that point of view, our data show that GSDMA and GSDMC were cleaved by caspase 3 during apoptosis (Fig. 2). Similarly, we found that GSDMD could be cleaved by caspase 3, and DFNA5 was cleaved by another caspase to enhance cell death and release ApoExos (Fig. 2), although it had previously been shown to be cleaved only by caspase 3 [9]. Thus, our results suggest that various caspases or enzymes could cleave gasdermins.

According to the findings of this study, the gasdermin-mediated promotion of ApoExo biogenesis somewhat resembles that membrane repair process in its mechanisms. The gasdermin-mediated process has been shown to depend on cytosolic Ca^2+^ and the ESCRT-III complex, and it requires annexins, PEF family proteins, sorcin/grancalcin, and HD-PTP. Accordingly, annexins might confer a platform for gathering sorcin/grancalcin-HD-PTP and eventually ESCRT-III during the ILV formation stage of ApoExo biogenesis. That mechanism agrees with recent reports showing that either sorcin or grancalcin can bind to annexins [30, 31, 32]. Consistent with our findings, GSDMD has been reported to promote plasma membrane repair by recruiting the ESCRT-III complex, which prolongs the survival of a cell dying through pyroptosis [33]. Our data are also supported by the novel role of MLKL (mixed lineage kinase domain-like protein), a pore-forming protein in necroptosis, which regulates membrane repair [34] and promotes exosome biogenesis using Ca^2+^-mediated ESCRT-III machinery [35].

Recent research has highlighted that gasdermins can affect the transition between apoptosis and pyroptosis [36]. Thus, it is noteworthy that immunologically, apoptosis can no longer be considered a quiescent type of cell death because from the beginning of apoptotic signaling, stressed cells or tissues can change their destinies to pyroptotic cell death by activating the gasdermins and thereby releasing their intracellular contents to adjacent cells to provoke inflammation and immune responses. As demonstrated in this study, the role of gasdermins in cell death is not confined to the transition between modes of death because the gasdermins can control the release of exosomes. Given that ApoExos are probably loaded with the contents of dying cells [37], they transfer information from dying cells to near or distant living cells and tissues, causing the regeneration of damaged tissues, pro- or anti-inflammatory effects, or the arousal of immune responses to autoantigens or cancer antigens [11, 12]. Therefore, a complete molecular understanding of the biogenetic mechanisms of ApoExos will be needed to comprehend the pathophysiologic events associated with apoptosis, which can then be used to develop novel therapeutic approaches.

## Materials and Methods

### Cells, antibodies, and other reagents

HeLa cells (human cervical cancer) were cultured in minimal essential medium supplemented with 10% heat-inactivated FBS, 2mM L-glutamine, 100U/ml penicillin, and 100 μg/ml streptomycin. MCF7 (human breast cancer), U251 (human malignant glioblastoma), HEK 293T (human embryonic kidney), HepG2 (human hepatocellular cancer), and 4T1 (mouse breast cancer) cells were cultured in DMEM supplemented with 10% heat-inactivated FBS, 2 mM L-glutamine, 100 U/ml penicillin, and 100 μg/ml streptomycin. A549 (human lung cancer) cells were cultured in 10% FBS RPMI1640. Staurosporine, cycloheximide, and BAPTA-AM were purchased from Santa Cruz Biotechnology (Santa Cruz, CA). TNFα was purchased from Sino Biological (Beijing, China). Alexa Fluor 594–conjugated wheat germ agglutinin, Alexa Fluor 594–conjugated anti-mouse IgG Ab, and Alexa Fluor 594–conjugated anti-rabbit IgG Ab were purchased from Thermo Fisher Scientific (Seoul, Korea). A23187 was purchased from Sigma-Aldrich (Yongin, Korea). Anti-GSDMA and anti-GSDMB Abs were purchased from ProSci (Fort Collins, CO); anti-GSDMC, anti-GSDMD, and anti-DFNA5 Abs were from Proteintech (Rosemont, IL); anti-DFNB59 and anti-peflin Abs were from Novus Biologicals (Centennial, CO); anti-CD63, anti-S1PR3, anti-V5 Tag, anti-GAPDH, anti-ANXA7, anti-GFP, and anti-grancalcin Abs were from Santa Cruz Biotechnology; anti-S1PR1 Ab was from Merck Millipore (Seoul, Korea); anti-Flag M2 Ab, anti-Flag M2 affinity gel, and anti-V5 affinity gel were from Sigma Aldrich (Yongin, Korea); anti-ANXA2 Ab was from Cell Signaling Technology (Danvers, MA); anti-sorcin and anti-HD-PTP Abs were from Thermo Fisher Scientific (Seoul, Korea); and anti-CHMP4B Ab was from GeneTex (Hsinchu City, Taiwan).

### Expression constructs and lentiviral transfection

CD63-pEGFP C2 was a gift from Paul Luzio (Addgene plasmid # 62964), and mCherry2-N1 was a gift from Michael Davidson (Addgene plasmid # 54517). Plasmids containing cDNAs of GSDMA, GSDMB, GSDMC, GSDMD, DFNA5, and DFNB59 were purchased from Sino-Biological (Beijing, China). The cDNAs of VPS4B, CHMP2A, CHMP3, ANXA2, ANXA7, ALG2, PEF1, sorcin, grancalcin, and HD-PTP were amplified from a mixture of HeLa cell cDNA. Dfna5 cDNA was amplified from a mixture of 4T1 cDNA. Constructs of cDNAs were cloned into pCDH-CMV-MCS-EF1-Puro, pCDH-CMV-MCS-EF1-Blast, pCDH-EF1-MCS-T2A-Puro, or pCDH-EF1-MCS-T2A-Blast with a C-terminal Flag tag, C-terminal V5 tag, or N-terminal Flag tag. Site-directed mutagenesis of GSDMD and DFNA5 was performed using a Quick-change site-directed mutagenesis kit (Thermo Fisher Scientific). For construction of DFNA5 fusion cDNA, a BsaBI site was introduced into the linker domain sequence of DFNA5 located N-terminally from the D270 caspase-cleavage site by site-directed mutagenesis, and Flag oligonucleotides, eGFP cDNA, or mCherry cDNA were cloned at the position next to S252 of DFNA5 cDNA using an In-Fusion HD cloning kit (TaKaRa, Seoul, Korea). All the lentiviral vectors, pGagPol (Sigma Aldrich), and pVSVg (Sigma Aldrich) were transfected into 293TN cells (System Biosciences) using Lipofectamine 2000 transfection reagent (Thermo Fisher Scientific). Particles were collected on the 2^nd^ and 3^rd^ days after the transfection of the lentiviral plasmids and infected into the cells. Lentivirus-infected cells were puromycin- or blasticidin-selected for 2 weeks.

### Cumate-inducible system for DFNA5 expression

Complete or partial DFNA5 cDNA was cloned into PiggyBac Cumate Switch Inducible vectors (PB-Cuo-MCS-IRES-GFP-EF1-CymR-T2A-Puro or PB-Cuo-MCS-3XFlag-IRES-GFP-EF1-CymR-T2A-Puro) (System Biosciences, Palo Alto, CA). The vectors were transduced into HeLa cells using the PiggyBac transposon system (System Biosciences, Palo Alto, CA), and the cells were selected for 2 weeks with 2 μg/ml of puromycin. For induction of DFNA5 cDNA, the cells were incubated with cumate (30 μg / ml) for 48 hr, and the expression of DFNA5 was confirmed by western blotting.

### CRISPR/Cas9 knockout and stable cell line generation

The generation of stable knockout cell lines was achieved using the LentiCRISPRv2 system (one-vector system, Addgene #52961) or LentiGuide-Puro system (two-vector system, Addgene #52963 and #52962), which were gifts from Feng Zhang. A lentiviral plasmid containing guide RNA sequences was transfected into 293TN cells (System Biosciences) using Lipofectamine 2000 transfection reagent. Particles were collected 2 days after transfection of the lentiviral plasmids and infected into HeLa cells. The lentivirus-infected HeLa cells were puromycin-selected for 2 weeks and tested by western blotting to confirm gene knockout. Cells showing decreased or knocked-out expression of the target genes were used in further experiments.

### Nanoparticle tracking analysis

Conditioned media from cell culture supernatants or plasma from EDTA-treated whole mouse blood were diluted tenfold with PBS and then analyzed for the number and size of particles using NTAs performed with a ZetaView Nanoparticle Tracking Analyzer. The following tracking parameters were used: camera sensitivity (85), shutter (250), frame rate (30f/s), minimum brightness (30), maximum area (1,000), minimum area (10), and traces (12).

### LDH assay

To detect pyroptosis among apoptotic cells, we performed the lactate dehydrogenase (LDH) assay according to the instructions of the manufacturer (Biovision, Milpitas, CA), with minor modification. Briefly, the cells were cultured in 24-well plates to more than 90% confluency and then treated with staurosporine (1 μM) or TNFα (50 ng/ml) and cycloheximide (5 μg/ml) for 6 to 48 hr. The culture supernatants were collected by centrifugation of the conditioned media for 10 min at 600Xg. Cell lysates were taken from the cellular pellets after centrifugation of the conditioned media, and adherent cells were taken by lysis with 1% Triton X100 PBS. The LDH assay was performed by mixing 100 μl of the diluted supernatants or cell lysate with 100 μl of reconstituted catalyst solution in clear 96-well plates in triplicate, incubating them for 30 min at room temperature, and then measuring the absorbance at 490 nm. % cell death was calculated as follows: (absorbance of supernatant) / (absorbance of cell lysate + absorbance of supernatant) X 100.

### Preparation of exosomal fractions

The conditioned media from apoptotic cells or mouse plasma diluted tenfold with PBS were centrifuged for 10 min at 200Xg and for 20 min at 2,000Xg twice to remove cellular debris and apoptotic bodies. Then the pellets were collected and washed by ultracentrifugation at 100,000Xg for 70 min twice. For fractionation of the vesicular membrane, the exosomes were incubated on ice with 100mM Na_2_CO_3_ (pH 11) for 1 hr, washed once, and resuspended in PBS.

### Confocal microscopy

Cells grown on Lab-Tek four-well glass chamber slides (NUNC), were incubated in medium or medium containing staurosporine (1 μM) for the indicated times. In some experiments, the cells were incubated with Alexa Fluor 594–conjugated wheat germ agglutinin (2.5 μg/ml) for 10 min at 37 °C and washed twice with HBSS. The cells were fixed with 4% paraformaldehyde and permeabilized with 0.2% Triton X100 PBS or fixed with methanol. The fixed cells were stained with the appropriate Ab and mounted with DAPI-containing mounting medium (Vector Laboratories Ltd, Peterborough, UK). Images were collected using a laser scanning confocal microscope LSM710 (Carl Zeiss, Oberkochen, Germany) equipped with argon (488 nm) and krypton (568 nm) lasers and using a x40 water immersion objective. Images were processed with ZEN 2009 light edition (Carl Zeiss).

### Transmission electron microscopy (TEM) and immunogold labeling

Cells were pelleted and washed twice with PBS. Fixation was performed with phosphate buffer (pH 7.4) containing 2.5% glutaraldehyde for 30 min at 4 °C. The pellets were rinsed twice with cold PBS, post-fixed in buffered OsO_4_, dehydrated in graded acetone, and embedded in Durcupan ACM resin (Fluka, Yongin, Korea). Ultrathin sections were obtained, mounted in copper grids, and counterstained with uranyl acetate and lead citrate. The specimens were observed with a Hitachi H-7600 TEM (Schaumburg, IL, USA) at 80 kV. The exosomes were resuspended in PBS, deposited onto formvar carbon-coated nickel grids for 60 min, washed with PBS, and fixed with 2% paraformaldehyde for 10 min. The exosome-coated grids were washed with PBS, transferred to a drop of the antibody, and incubated for 40 min for the immunogold labeling with anti-DFNA5 antibody. The grids were washed with 0.1% BSA/PBS, incubated with 10 nm-gold labeled goat-anti-rabbit IgG for 40 min, washed in PBS, and then post-fixed with 2.5% glutaraldehyde for 10 min. After washing in deionized water, the grids were stained with 2% uranyl acetate for 15 min and with 0.13% uranyl acetate and 0.4% methylcellulose for 10 min, air-dried for 5 min, and viewed by TEM.

### Flow-cytometric analysis of exosomes

20 μg of exosomes were coated onto 5 μl of aldehyde/sulfate latex beads (4 μm in diameter) for 15 min at room temperature in PBS, with a final volume of 20 μl. The beads were then washed with 1 ml of PBS with shaking for 1 hr, blocked by incubation with 20 μl of FBS for 30 min, and washed thrice in PBS. The beads were resuspended in 50 μl of PBS and incubated with anti-DFNA5 Ab or isotype-matched irrelevant Ab for 1 hr at room temperature. After being washed thrice with PBS, the beads were incubated with PE-conjugated secondary Ab (Santa Cruz Biotechnology, Santa Cruz, CA) for 1 hr at room temperature. Finally, the beads were analyzed by flow cytometry using a FACSCalibur flow cytometer (Becton Dickinson, Mountain View, CA) and FlowJo software.

### Preparation of cell lysates and western blots

To prepare the lysates, cells and exosome pellets were lysed in lysis buffer (50 mM Tris-Cl, pH 7.5, 150 mM NaCl, 1 mM EDTA, 1% Triton X100, 1 mM Na_3_VO_4_, 1 mM NaF, 1 μg/ml pepstatin A, 10 μg/ml AEBSF, 2 μg/ml aprotinin, and 1 μg/ml leupeptin), incubated on ice for 20 min, and centrifuged for 20 min to remove the supernatants. The lysates were subjected to SDS-PAGE. The proteins were then electro-transferred to PVDF membranes and incubated overnight with antibodies at 4 °C. Subsequently, the membranes were incubated with peroxidase-conjugated secondary antibodies (Pierce, Rockford, IL, USA) for 1 h at room temperature, and the signal was detected using an enhanced chemiluminescence detection kit (Amersham Biosciences, Seongnam, Korea).

### Immunoprecipitation

Cells (1 × 10^7^ cells) were lysed using 1 ml of lysis buffer (250 mM sucrose, containing 10 mM HEPES, pH 7.4, 0.1% Triton X-100, 1 mM Na_3_VO_4_, 1 mM NaF, 1 μg/ml pepstatin A, 10 μg/ml AEBSF, 2 μg/ml aprotinin, and 1 μg/ml leupeptin) with and without 100 μM CaCl_2_ for 30 min at 4 °C and then centrifuged for 20 min at 13,000 rpm at 4 °C. The supernatants were stored at −070 °C. The cell lysates were precleared with protein A/G-Agarose (Santa Cruz Biotechnology, Santa Cruz, CA) by incubation for 1 h at 4 °C with constant agitation. The precleared lysates were then incubated for overnight with anti-Flag M2 affinity agarose or anti-V5 affinity agarose at 4 °C. The immunoprecipitates were washed six times in PBS. An aliquot of each sample was subjected to western blot analysis.

### 4T1 orthotropic breast cancer model

All animal experiments were performed following guidelines of and protocols approved by the *Laboratory Animal Research Center of Ajou University Medical Center*, Suwon, South Korea. Female BALB/c mice (8 weeks old) weighing 25 g were purchased from OrientBio (Seongnam, Korea). 4T1 cells (2 X 10^6^ cells) were injected into their mammary fat pads. Around 3 weeks after tumor injection, when the tumors had reached 1,000 mm^3^, the mice were injected intraperitoneally with PBS or cyclophosphamide monohydrate (BioVision, Milpitas, CA) (50 mg / kg) and doxorubicin hydrochloride (Santa Cruz Biotechnology) (3 mg / kg) twice with an interval of 24 hr. 24 hr after the last injection, the animals were sacrificed, and their livers, flushed with ice-cold PBS, and spleens were resected. The whole blood from the mice was collected by heart puncture, and the plasma was separated.

### Real-time PCR

Total RNA was isolated using an RNeasy kit (Qiagen, Seoul, Korea). A PrimeScript RT reagent kit (TaKaRa, Seoul, Korea) was used to reverse-transcribe mRNA into cDNA. PCR was then performed on a QuantStudio 3 machine (Thermo Fisher Scientific) using SYBR Premix Ex Taq II (TaKaRa). The analysis of each sample in triplicate was performed more than twice for each experiment, and data in the figures are reported as relative quantification: average values of 2^-ΔΔCT^±S.D.

### Measurements of intracellular free calcium ions

Intracellular free Ca^2+^ was measured using Calcium Sensor Dye eFluor^TM^ 514 (eBiosciences, San Diego, CA, USA). Briefly, cells were incubated for 30 min at 37 °C in medium containing 10 μM eFluor^TM^ 514 and then washed twice. Fluorescence was measured at an excitation wavelength of 490 nm and an emission wavelength of 514 nm with a FLUOstar Optima Microplate Fluorometer (BMG Labtech, Cary, NC, USA). Data are presented as relative fluorescence: MFI of treated cells / MFI of non-treated cells.

### Subcellular fractionation by differential centrifugation

Preparation of the LMF and HMF was performed as described previously [38] with modifications. HeLa cells were homogenized in sucrose buffer (250 mM sucrose, 10 mM HEPES, pH 7.4, 0.1 mM CaCl_2_) with protease inhibitors (1 mM Na_3_VO_4_, 1 mM NaF, 1 μg/ml pepstatin A, 10 μg/ml AEBSF, 2 μg/ml aprotinin, and 1 μg/ml leupeptin) by passing them through a 21-gauge needle. The homogenates were centrifuged at 1,200 X g for 5 min to pellet the nuclear fraction and broken cellular debris. The supernatants were further centrifuged at 15,000 X g for 10 min to separate the HMF and post-nuclear supernatants. The post-nuclear supernatants were then centrifuged for 1.5 h at 130,000 X g to isolate the LMF and cytosolic fraction.

### Statistical analyses

The values are presented as the mean ± S.D. Statistical analyses were performed using one-way analysis of variance (ANOVA) with either all pairwise multiple comparisons or multiple comparisons versus control (Holm-Sidak method). Statistical significance was considered at the P < 0.05 level.

## Author Contribution Statement

J.H., Y.J.K., D.A.C., D.W.K., and J.K. performed most of experiments. H.S.Y. performed the animal studies. S.A.S. performed the transmission EM, the flow cytometry and the DFNB59 experiments. T.M. and D.Y.K. wrote the paper with assistance from all authors. Y.J.C. designed the experimental concept and the individual experimental plans, and wrote the paper.

## Availability of Data and Materials

All data needed to evaluate the conclusions in the paper are present in the paper and/or the Supplementary Materials. Further information and requests for resources and reagents should be directed to the corresponding author, Y.J.C.

