## Supplementary Information for "Role of Gasdermins in the Biogenesis of Apoptotic Cell–Derived Exosomes"

**Supplementary Figures and Legends**

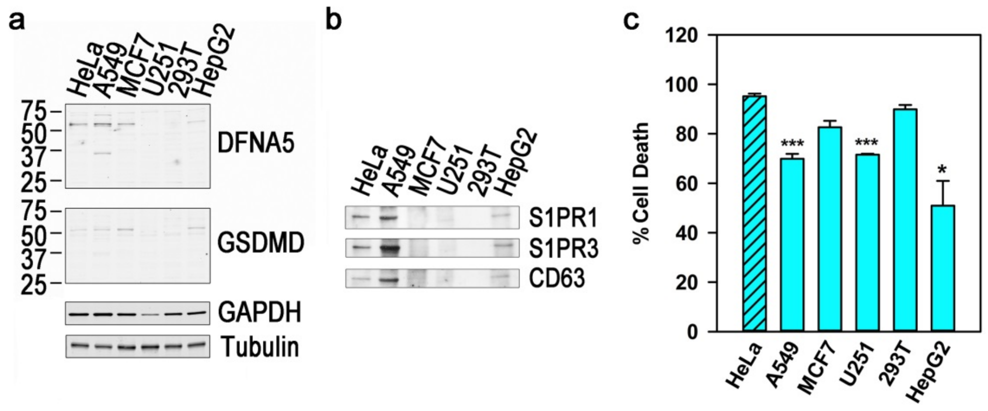


**Figure S1. Comparison between the expression of DFNA5 and GSDMD and the release of apoptotic exosomes in various cell lines.** (**a**) The expression of DFNA5 and GSDMD was inspected by western blotting. (**b**) The cells were treated with TNFα (50 ng/ml) and cycloheximide (25 μg/ml) for 48 hr. The exosomal fractions were prepared from the conditioned media and lysed with lysis buffer, with the volume adjusted for the cellular protein masses. Equal volumes of the exosomal fraction were western-blotted for the markers of apoptotic exosomes (*CD63*, *S1PR1*, and *S1PR3*). (**c**) % cell death was measured by the LDH assay, *P < 0.001, and ***P < 0.05.

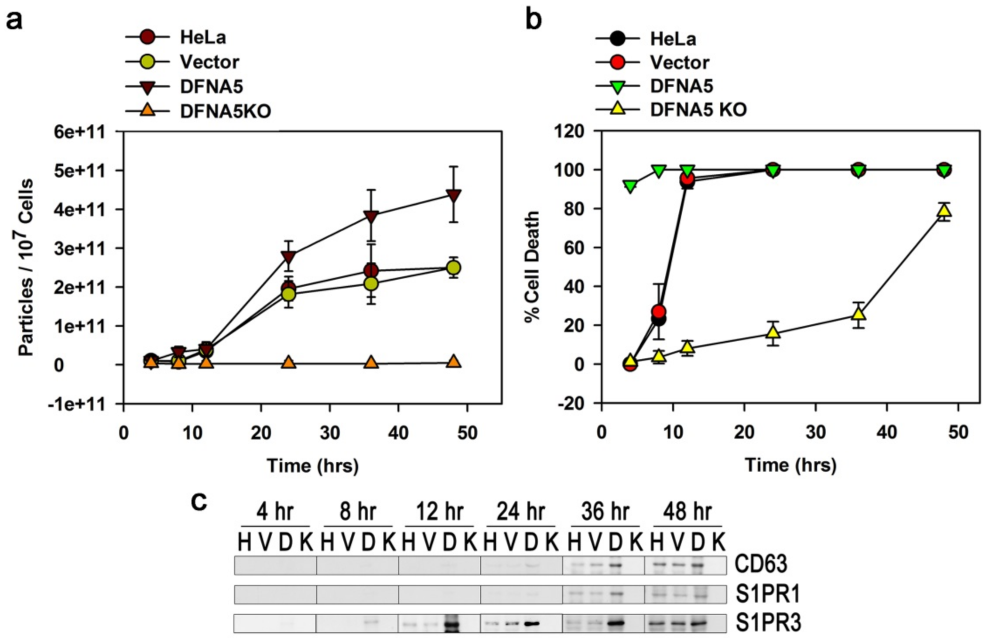


**Figure S2. DFNA5 is a prerequisite for the biogenesis of apoptotic exosomes.** Parental HeLa cells, cells infected with a control vector (*vector*), cells overexpressing DFNA5 (*DFNA5*), and DFNA5 knockout cells (*DFNA5KO*) were incubated with staurosporine (1 μM) for the indicated times. (**a**) Conditioned media were prepared, and the number of apoptotic exosomes was measured by nanoparticle-tracking analysis (NTA). (**b**) Pyroptotic cell death was detected by the LDH assay. (**c**) The exosomal fractions were prepared and western-blotted for the markers of apoptotic exosomes (CD63, S1PR1, and S1PR3) (*H*: from parental HeLa cells; *V*: vector-infected HeLa cells; *D*: cells overexpressing DFNA5; *K*: DFNA5 knockout cells).

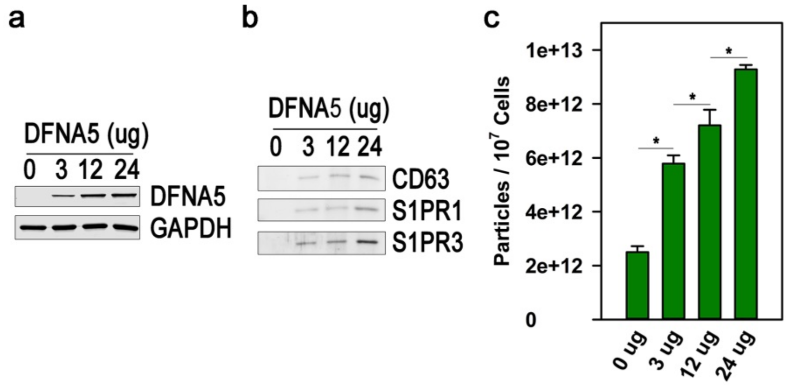


**Figure S3. DFNA5 enhances the release of apoptotic exosomes in a dose-dependent manner.** (**a**) HeLa cells grown to 90% confluency in a 100 mm dish were transfected with a plasmid containing DFNA5 cDNA in the indicated amounts. 24 hr after transfection, the cells were collected, and cytoplasmic protein was extracted. DFNA5 expression was observed by western blotting. (**b**) The cells were treated with staurosporine (1 μM) for an additional 24 hr, and apoptotic exosomes were prepared from the conditioned media. The markers of apoptotic exosomes (*CD63, S1PR1,* and *S1PR3*) were detected by western blotting from equal volumes of apoptotic exosomes. (**c**) Exosomal fractions were analyzed by NTA. * $P < 0.001$.

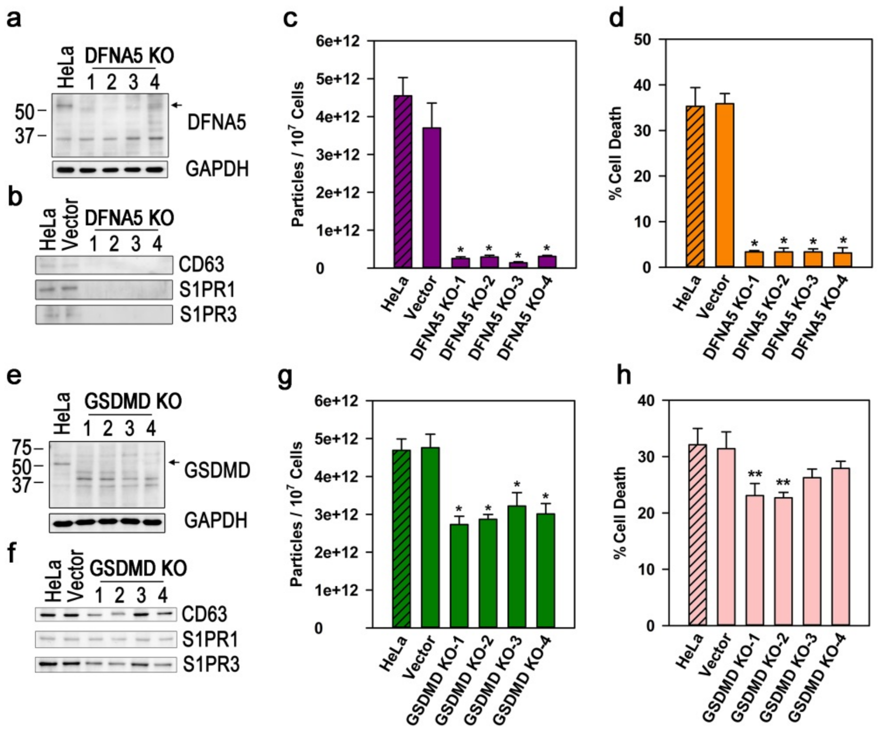


**Figure S4. Knockout of either DFNA5 or GSDMD hinders the release of apoptotic exosomes.** (**a** and **e**) Either DFNA5 or GSDMD was knocked out in HeLa cells by the CRISPR/Cas9 system. Arrows indicate the full-length forms of DFNA5 and GSDMD. (**b** and **f**) The cells were treated with staurosporine (1 μM). 24 hr after treatment, the apoptotic exosomes were purified from the conditioned media, western-blotted for CD63, S1PR1, and S1PR3. (**c** and **g**) The exosomes were analyzed by NTA. (**d** and **h**) Pyroptotic cell death was measured by the LDH assay. * $P < 0.001$ and ** $P < 0.01$ (**c, d, g,** and **h**).

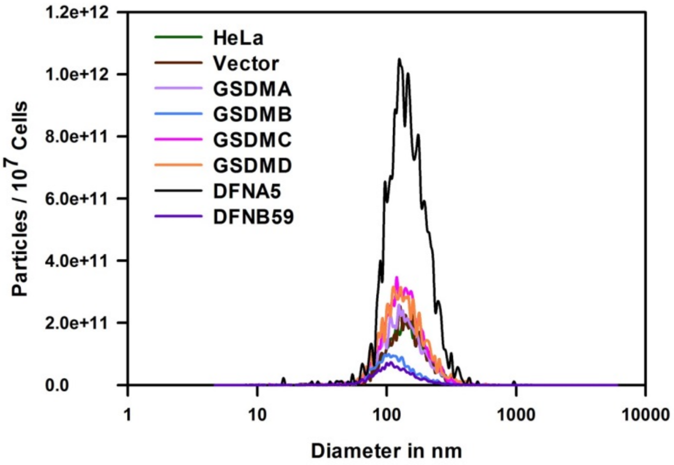


**Figure S5. Overexpression of GSDMA, GSDMC, GSDMD, or DFNA5 increases the release of apoptotic exosomes.** HeLa cells overexpressing gasdermins were treated with staurosporine (1 μM) for 24 hr. Exosomal fractions were prepared from the conditioned media and analyzed by NTA.

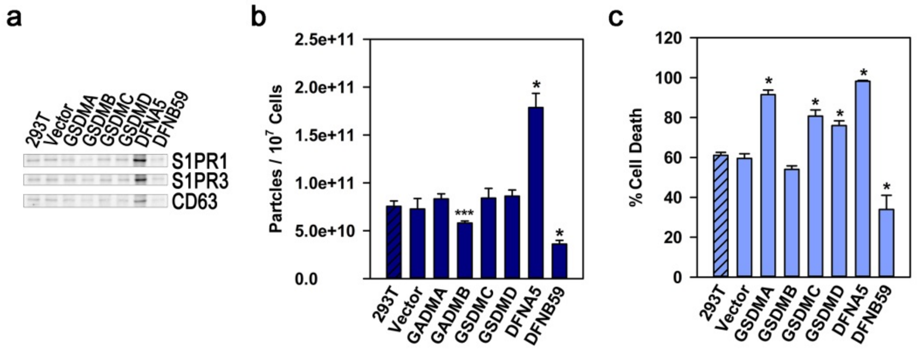


**Figure S6. Gasdermins regulate the release of apoptotic exosomes in staurosporine-treated 293T cells.** (**a** and **b**) Parental 293T cells and 293T cells infected with a control vector or a gasdermin (*GSDMA*, *GSDMB*, *GSDMC*, *GSDMD*, *DFNA5*, or *DFNB59*) were incubated with staurosporine (10 μM). 48 hr after treatment, exosomal fractions were prepared from the conditioned media, and equal volumes of apoptotic exosomes were western-blotted for markers of apoptotic exosomes (*S1PR1*, *S1PR3*, or *CD63*) and analyzed by NTA. (**c**) Pyroptotic cell death was measured by the LDH assay 24 hr after staurosporine treatment. * P < 0.001 and *** P < 0.05 (**b** and **c**).

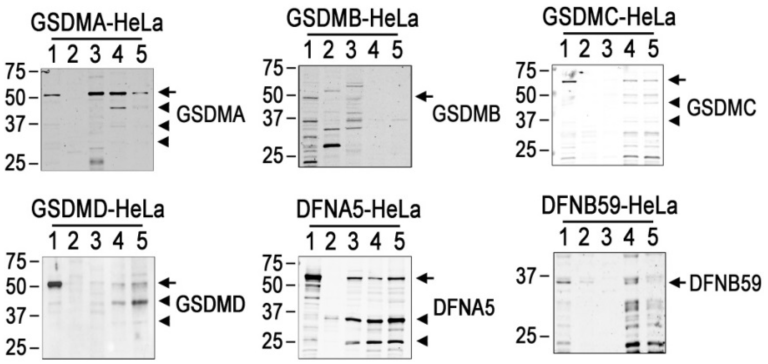


**Figure S7. GSDMA, GSDMC, GSDMD, and DFNA5 are localized in the membrane of apoptotic exosomes.** HeLa cells overexpressing gasdermins were treated with staurosporine (1 μM) for 24 hr. Equal amounts of protein extracted from non-treated control cells, staurosporine-treated cells, conditioned media, apoptotic exosomes, and membrane fractions of apoptotic exosomes were western-blotted for the indicated gasdermins. Arrows denote the full-length form of each gasdermin, and triangles denote the cleaved forms (*1*: non-treated control cells; *2*: staurosporine-treated cells; *3*: conditioned media; *4*: apoptotic exosomes; *5*: membrane fractions of apoptotic exosomes).

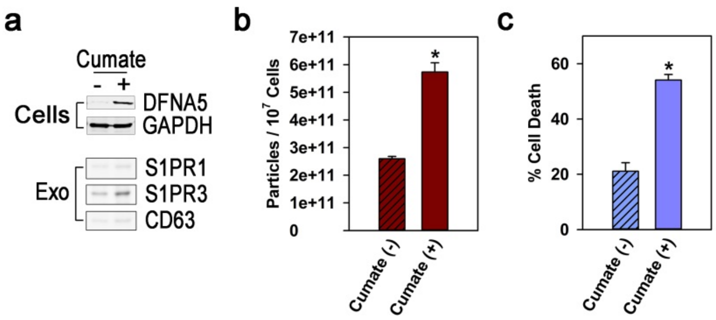


**Figure S8. Full-length DFNA5 induced by cumate treatment increases the release of ApoExos and early pyroptotic cell death.** (**a, upper panel**) Full-length DFNA5 cDNA under the control of a cumate-inducible promotor was stably expressed in HeLa cells. The cells were treated with or without cumate (20 μg/ml) for 48 hr. Induction of DFNA5 was confirmed by western blotting. (**a, lower panel** and **b**) Apoptotic cell death was guided by staurosporine treatment (1 μM) for 24 hr. ApoExos were prepared from the conditioned media and analyzed for ApoExo markers by western blotting and NTAs. (**c**) Pyroptotic cell death was measured by the LDH assay. *P < 0.001 (**b** and **c**).

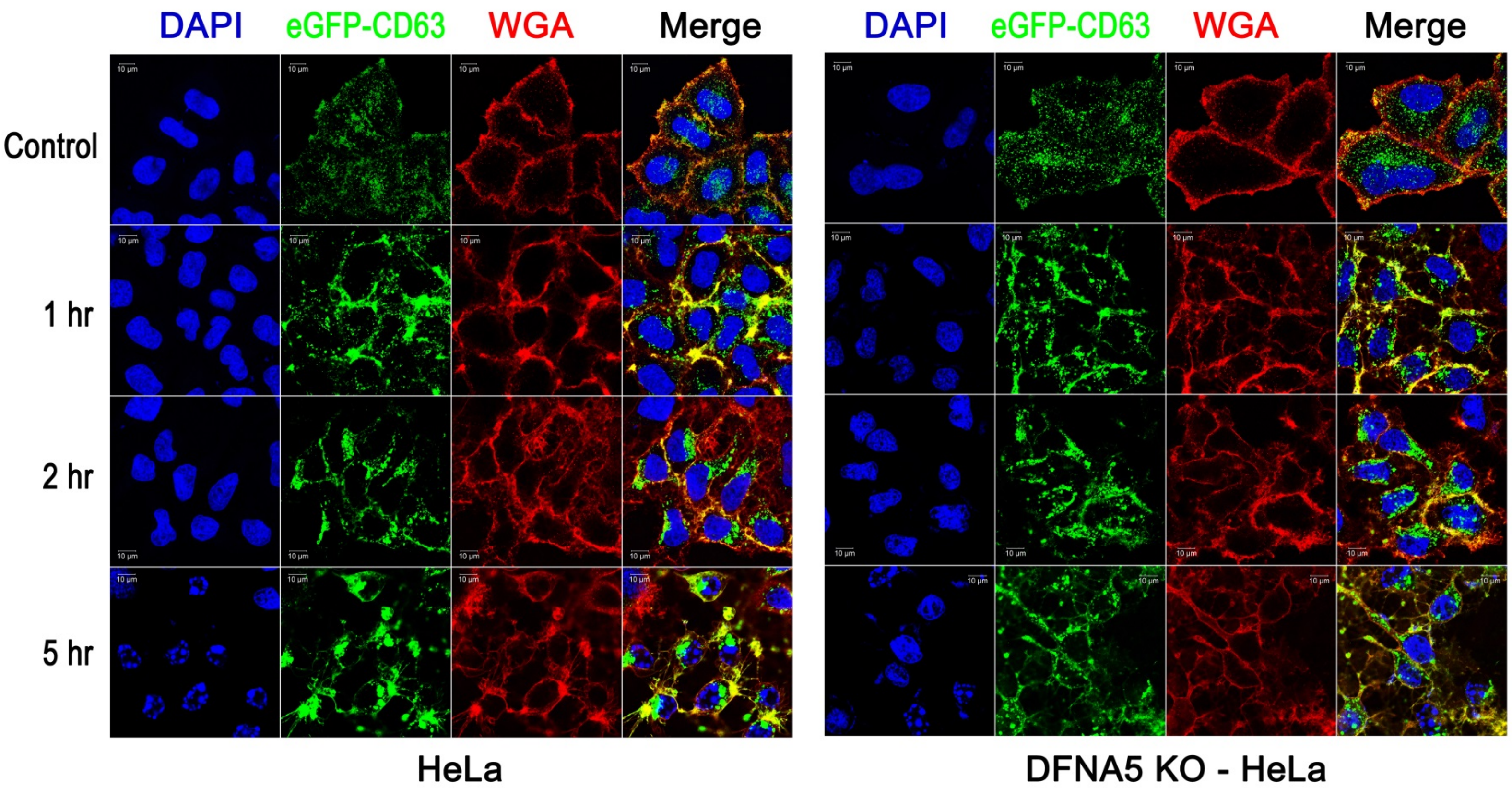


**Figure S9. DFNA5 KO diminished the large intracellular vesicular aggregates that appear in apoptotic cells.** Parental HeLa cells and DFNA5 knockout cells expressing eGFP-CD63 were treated with staurosporine (1 μM) for the indicated times. The cells were stained with wheat germ agglutinin-Alexa Fluor 594 (*WGA*), and confocal images are shown.

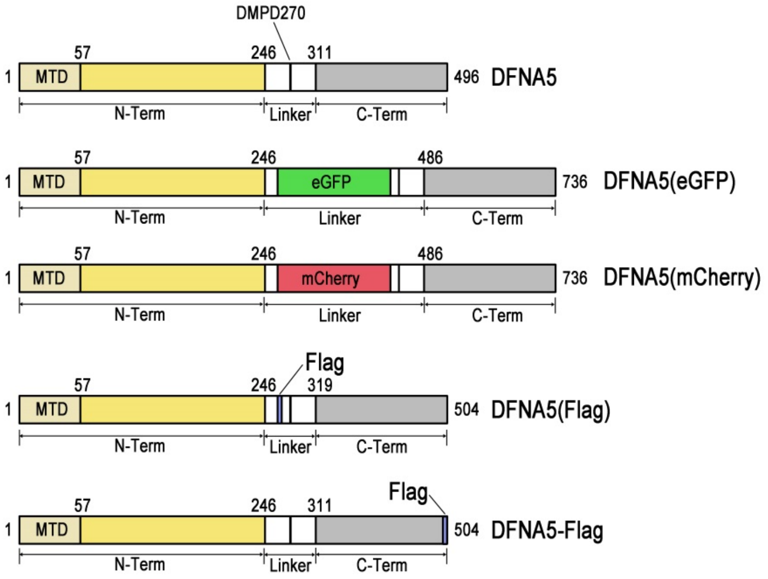


**Figure S10. Schematic diagram of DFNA5 constructs.** DFNA5 is composed of an N-terminal membrane translocation domain (*MTD*), pore-forming N-terminal segment (*N-Term*), C-terminal segment (*C-Term*), and linker region connecting the N-terminal and C-terminal segments (*Linker*). eGFP, mCherry, or Flag tag sequences were inserted into the linker region amino-terminally from the caspase-cleavage site (Asp270) [*DFNA5(eGFP), DFNA5(mCherry),* and *DFNA5(Flag)*]. The Flag tag was added at the C-terminal end of DFNA5 (*DFNA5-Flag*). The asterisk indicates the caspase-cleavage site of DFNA5.

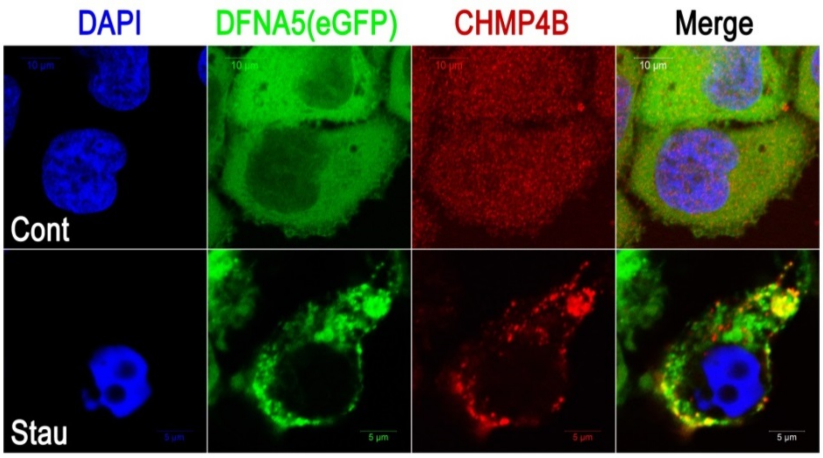


**Figure S11. CHMP4B is co-localized with DFNA5 in apoptotic cells.** HeLa cells expressing DFNA5 with internal eGFP were incubated with solvent or staurosporine (1 μM) for 4 hr and then stained for CHMP4B. Confocal images of the cells are shown.

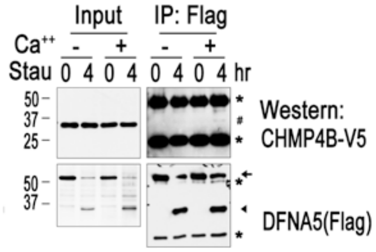


**Figure S12. Interaction between DFNA5 and CHMP4B depends on $Ca^{2+}$.** HeLa cells expressing DFNA5 with an internal Flag tag [DFNA5(Flag)] and CHMP4B-V5 were treated with staurosporine (1 μM) for the indicated times. Cellular lysates were prepared with and without $Ca^{++}$ (100 μM), of which one tenth of the volume was western-blotted for V5 and Flag (*Input*). The immunoprecipitates with anti-Flag Ab were western-blotted with anti-Flag and anti-V5 Abs (*IP: Flag*). The arrow and triangle denote full-length and cleaved N-terminal segments of DFNA5, respectively. Asterisks indicate Ig heavy and light chains, and # indicates co-immunoprecipitated CHMP4B-V5.

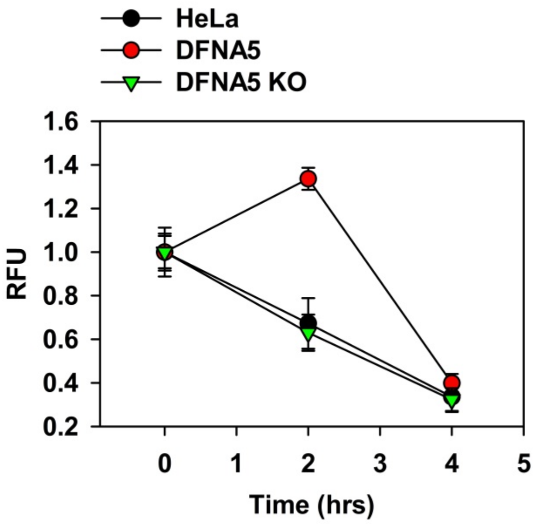


**Figure S13. Overexpression of DFNA5 transiently increases intracellular free $Ca^{2+}$.** Intracellular free $Ca^{2+}$ was measured with a calcium sensor dye, eFluor™ 514, at the indicated times in staurosporine-treated wild-type, DFNA5-overexpressing, and DFNA5-depleted HeLa cells. The data were analyzed as relative fluorescence units (*RFU*): fluorescence intensities of the staurosporine-treated cells / fluorescence intensities of the non-treated control cells. The data pictured are from one representative experiment executed in quintuplicate from among three independent experiments.

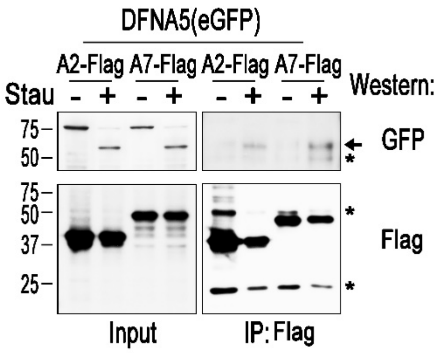


**Figure S14. DFNA5 is co-immunoprecipitated with Annexin A2 and A7 in apoptotic HeLa cells**. HeLa cells expressing ANXA2-Flag or ANXA7-Flag together with DFNA5(eGFP) were treated with staurosporine (1 µM) or DMSO for 4 hr. The cell lysates were immunoprecipitated with agarose beads conjugated with anti-Flag Ab. One hundredth of the cell lysates (*Input*) and immunoprecipitates (*IP: Flag*) were western-blotted for the Flag or GFP tag. Arrows indicate DFNA5(eGFP), and asterisks indicate Ig heavy and light chains.

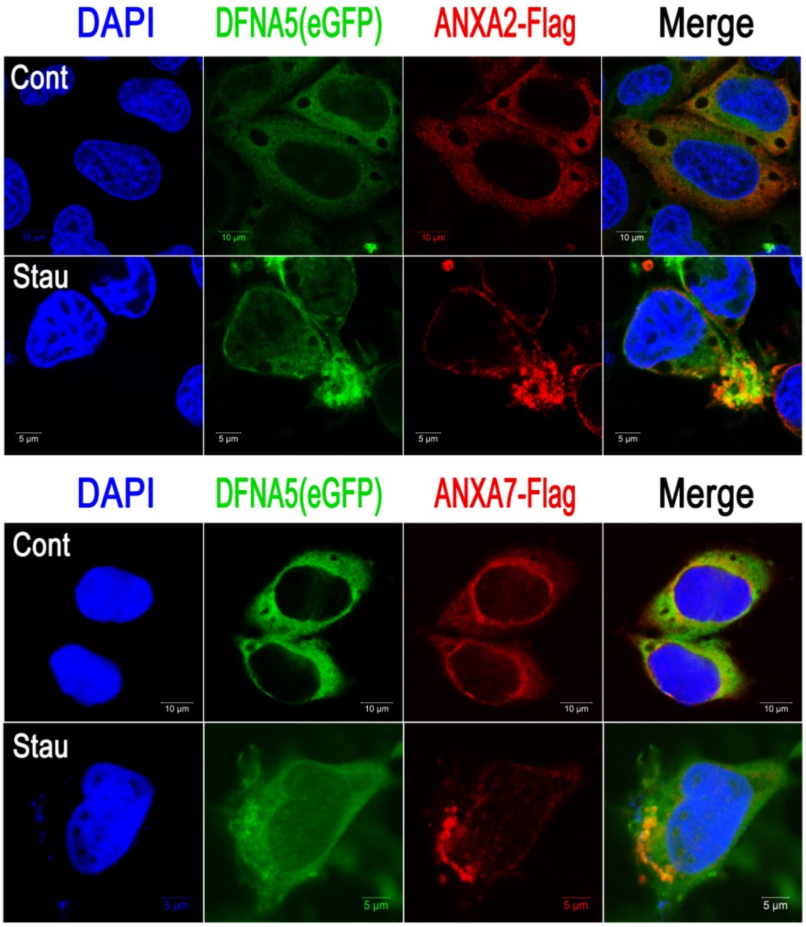


**Figure S15. Annexin A2 and annexin A7 are co-localized with DFNA5 in apoptotic cells.** HeLa cells expressing DFNA5(eGFP) with ANXA2-Flag or ANXA7-Flag were treated with staurosporine (1 μM) or control solvent (DMSO) for 4 hr and stained with anti-Flag Ab and Alexa Fluor 594–conjugated anti-mouse Ig Ab. Confocal images of the cells are shown.

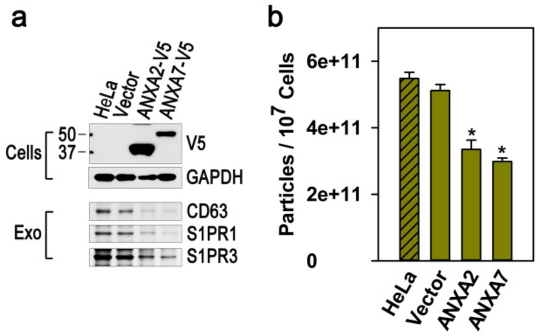


**Figure S16. Overexpression of ANXA2 or ANXA7 inhibits the release of apoptotic exosomes.** (**a, upper panel**) Control HeLa cells and cells overexpressing ANXA2 or ANXA7 tagged with V5 were western-blotted for V5-tag and GAPDH. (**a, lower panel** and **b**) The cells were treated with staurosporine (1 μM) for 24 hr. The exosomes were prepared and analyzed by western blotting or NTAs. *P < 0.001 (**b**).

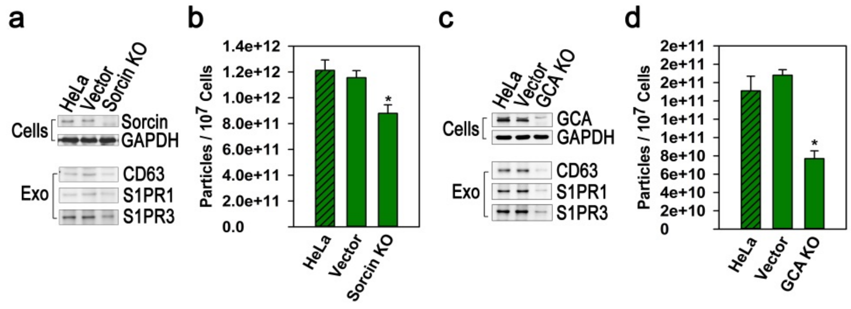


**Figure S17. Depletion of sorcin reduces the release of apoptotic exosomes. (a, upper panel** and **c, upper penal)** Control HeLa cells (*HeLa* and *Vector*) and cells depleted of the sorcin gene (*Sorcin KO*) or the grancalcin (*GCA KO*) were western-blotted for sorcin, grancalcin, or GAPDH. (**a, lower panel** and **c, lower panel**) The cells were treated with staurosporine (1 μM) for 24 hr. Apoptotic exosomes purified from the conditioned media were analyzed via western blotting. (**b** and **d**) The exosomes were analyzed by NTAs. *P < 0.001 (**b** and **d**).

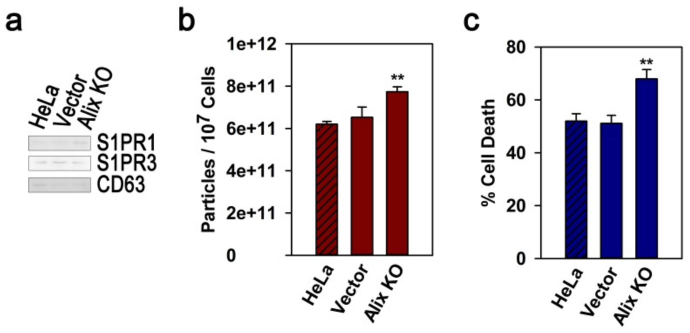


**Figure S18. Alix is not associated with the biogenesis of apoptotic exosomes.** (**a** and **b**) Control HeLa cells (*HeLa* and *Vector*) and Alix knockout cells (*Alix KO*) were treated with staurosporine (1 μM) for 24 hr. Apoptotic exosomes were prepared and analyzed by western blotting and NTAs. (**c**) Pyroptotic cell death was measured in cells treated with staurosporine for 12 hr. **P < 0.01 (**b** and **c**).

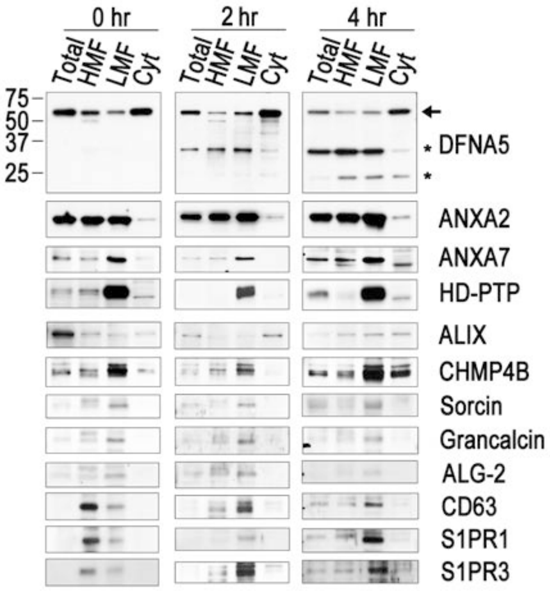


**Figure S19. DFNA5 migrated into MVB-containing LMFs is co-localized with ANXA2, ANXA7, sorcin, grancalcin, HD-PTP, and CHMP4B in apoptotic cells.** DFNA5-overexpressing HeLa cells were treated with staurosporine (1 μM) for the indicated times. The total cellular lysates (*Total*), heavy membrane fraction (*HMF*), light membrane fraction (*LMF*), and cytosolic fraction (*Cyt*) were then prepared. Equal amounts of protein were separated by SDS-PAGE and western-blotted for the indicated proteins. Arrows and asterisks denote the full-length and cleaved forms of DFNA5, respectively.

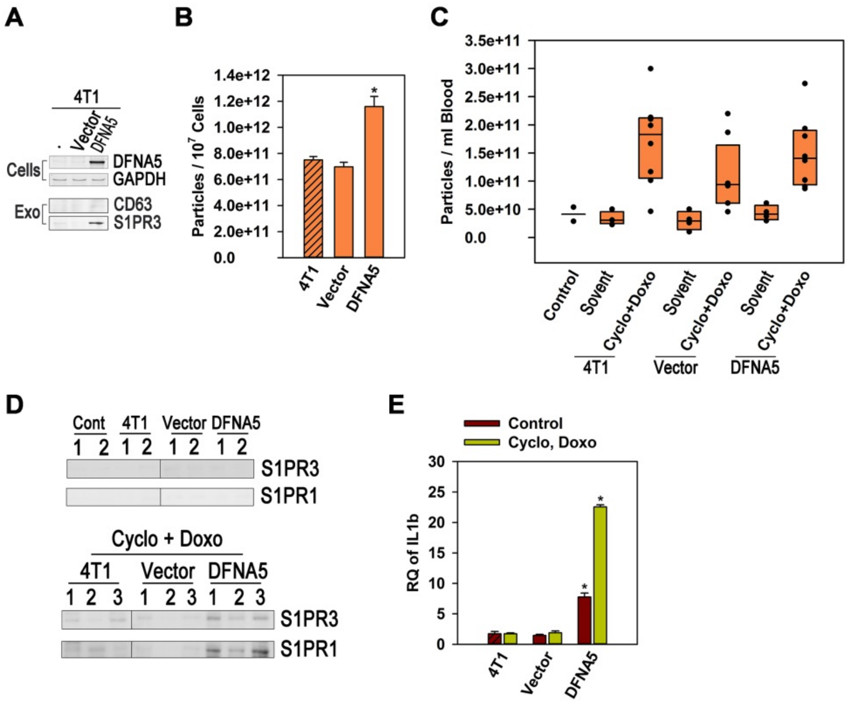


**Figure S20. DFNA5 overexpression enhances the level of plasma apoptotic exosomes and inflammation in the 4T1 orthotopic breast cancer model.** (**a, upper panel**) Parental 4T1 cells and cells infected with control vector or Dfna5 cDNA were western-blotted to affirm Dfna5 expression. (**a, lower panel** and **b**)The cells were treated with staurosporine (1 μM) for 24 hrs. The released ApoExos were measured by western blotting and NTAs. (**c** and **d)** BALB/c mice were injected in their mammary fat pads with 4T1, vector-infected 4T1, or Dfna5-infected 4T1 cells, and tumors were grown for around 3 weeks, until their mass reached 1,000 $mm^3$. Plasma exosomes were prepared from the mice intraperitoneally treated with solvent or cyclophosphamide (50 mg/kg) and doxorubicin (3 mg/kg) and analyzed by NTAs (**c**) and western blotting for the markers of ApoExos (**d**). (**e**) mRNA expression of inflammatory mediators in liver tissue was measured by real-time PCR. *P < 0.001 (**b,** and **e**).

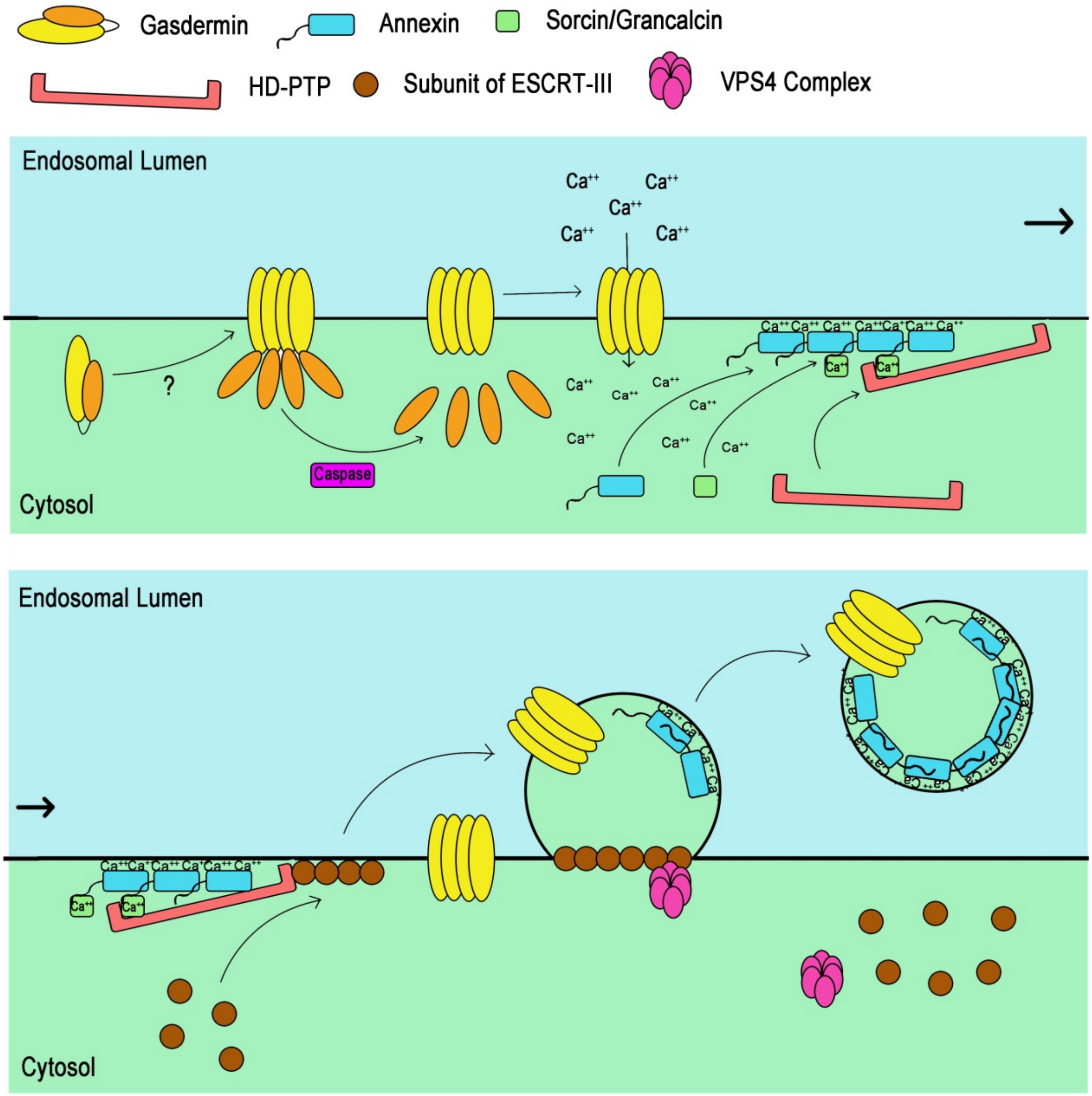


**Figure S21. Schematic illustration summarizing gasdermin-mediated biogenesis of apoptotic exosomes.** In the apoptotic cells, gasdermins migrate toward the endosomal membrane under the influence of unknown factors and are cleaved by caspases, thereby producing N-terminal fragments that form pores at the endosomes. $Ca^{2+}$ enters the cytosol through those pores, increasing the regional $Ca^{2+}$ concentration, which sequentially recruits $Ca^{2+}$-binding proteins (annexins), sorcin/grancalcin, and HD-PTP to the endosomal membrane. Members of the ESCRT-III complex bind to the Bro1 domain of HD-PTP and begin to make outward buds in the endosomal membrane. Eventually, the budded endosomal membrane is liberated as intraluminal vesicles (ILVs) by membrane fission with the help of the VPS4 complex, which contributes to the fission of the endosomal membrane and disassembly of the ESCRT-III complex for recycling. The luminal side of the ILV membrane is coated by annexins such as ANXA2, which possibly function as membrane scaffolds.
